# Frameshifting at collided ribosomes is modulated by elongation factor eEF3 and by Integrated Stress Response regulators Gcn1 and Gcn20

**DOI:** 10.1101/2021.08.26.457827

**Authors:** Lisa Houston, Evan Platten, Sara Connelly, Jiyu Wang, Elizabeth J. Grayhack

## Abstract

Ribosome stalls can result in ribosome collisions that elicit quality control responses, one function of which is to prevent frameshifting by the stalled ribosome, an activity that entails interaction of the conserved yeast protein Mbf1 with uS3 on colliding ribosomes. However, the full spectrum of factors that mediate frameshifting during ribosome collisions is unknown. To delineate such factors in the yeast *Saccharomyces cerevisiae*, we used genetic selections for mutants that either suppress or increase frameshifting from a known ribosome stall site, CGA codon repeats. We show that the general translation elongation factor eEF3 promotes frameshifting, while Integrated Stress Response (ISR) pathway components Gcn1 and Gcn20 suppress frameshifting. We found a mutant form of eEF3 that specifically suppressed frameshifting, but not translation inhibition by CGA codons. Thus, we infer that frameshifting at collided ribosomes requires eEF3, which facilitates tRNA-mRNA translocation and E-site tRNA release in yeast and other single cell organisms. By contrast, we found that removal of either Gcn1 or Gcn20, which bind collided ribosomes with Mbf1, increased frameshifting. Thus, we conclude that frameshifting is suppressed by Gcn1 and Gcn20, although these effects are not mediated through activation of the ISR. Furthermore, we examined the relationship of eEF3-mediated frameshifting to other quality control mechanisms, finding that the eEF3-mediated frameshifting competes with No-Go decay, Mbf1 and Gcn1/20. Thus, these results provide evidence of a direct link between translation elongation and frameshifting at collided ribosomes, as well as evidence that frameshifting competes with other quality control pathways that act on collided ribosomes.

## INTRODUCTION

Ribosomes not only accurately translate an open reading frame into the specified amino acid sequence, but also display the plasticity to accommodate regulatory events during elongation. To this end, ribosomes elongate the nascent chain at variable speeds, using high rates to maximize protein production, and lower rates and stalls to assist regulatory events, such as protein folding, localization, protein-protein interactions and programmed frameshifting (Collart and Weiss 2020). Ribosomes also stall during translation for a variety of reasons, including sequences and structures encoded in the mRNA, the composition of the nascent peptide, damage to the mRNA and stochastic events (Doma and Parker 2006; Dimitrova et al. 2009; Letzring et al. 2013; Simms et al. 2014; Brandman and Hegde 2016; Gamble et al. 2016; Joazeiro 2017).

Some ribosomes fail to efficiently resume translation after these stalls and must be resolved by means other than continued translation. Organisms in all kingdoms have developed quality control mechanisms that act on mRNAs with stalled ribosomes (Brandman et al. 2012; Samatova et al. 2020; D’Orazio and Green 2021). In some cases, ribosome stalls result in ribosome collisions that have been implicated as the trigger for quality control responses, which act to remove stalled ribosomes from the mRNA, degrade defective mRNAs and incomplete proteins, prevent loss of reading frame by the stalled ribosome, and activate global stress response pathways [see (Meydan and Guydosh 2021)].

Two non-ribosomal proteins conserved in eukaryotes regulate key local responses to ribosome stalls. Yeast Hel2 (ZNF598 in humans, designated hZNF598) (Garzia et al. 2017; Juszkiewicz et al. 2018; Ikeuchi et al. 2019) ubiquitinates 40S ribosomal proteins uS10 and uS3 (eS10, uS10, uS3 in humans) (Garzia et al. 2017; Juszkiewicz and Hegde 2017; Matsuo et al. 2017; Sundaramoorthy et al. 2017), promoting disassembly of ribosomal subunits of the lead ribosome by Slh1/Rqt2 (hASCC3 in humans) (Matsuo et al. 2017; Juszkiewicz et al. 2020b; Matsuo et al. 2020), recruitment of an endonuclease to the mRNA (D’Orazio et al. 2019; Glover et al. 2020), degradation of the mRNA by No-Go decay (NGD) (Doma and Parker 2006; Brandman et al. 2012; Saito et al. 2015; Brandman and Hegde 2016; Matsuo et al. 2017) and recognition of the released 60S subunit by the ribosome quality control (RQC) complex (Shao et al. 2013; Lyumkis et al. 2014; Shen et al. 2015). The RQC complex then targets the nascent peptide for degradation by ubiquitination (Brandman et al. 2012; Shao et al. 2013; Shao and Hegde 2014). The second factor, yeast Mbf1 (hEDF1 in humans) prevents the leading ribosome from frameshifting (Wang et al. 2018; Juszkiewicz et al. 2020a), although both the magnitude and directionality of frameshifting differ between yeast and humans. In humans, hEDF1 also promotes recruitment of hGIGYF2 and hEIF4E2, which in turn reduce translation initiation on mRNAs with collided ribosomes (Juszkiewicz et al. 2020a; Sinha et al. 2020). The yeast homologs of hGIGYF2, Smy2 and Syh1 are involved in decay of mRNAs with a stalling sequence (Hickey et al. 2020), but it is unknown how they are recruited to these mRNAs.

Induction of global stress-response pathways is also mediated through ribosome collisions. In both yeast and humans, ribosome collisions result in activation of the integrated stress response (ISR) kinase Gcn2, resulting in a global reduction in translation initiation (Meydan and Guydosh 2020; Wu et al. 2020; Pochopien et al. 2021; Yan and Zaher 2021). The observation that key effectors of the ISR, Gcn1, Gcn20, Rbg2 and Gir2, bind collided ribosomes with Mbf1 (Pochopien et al. 2021) likely explains a previous observation that Mbf1 also modulates the induction of the ISR in yeast (Takemaru et al. 1998). In humans, the ribosome-associated MAPKKK hZAKα autophosphorylates during ribosome collisions, resulting in activation of stress-activated protein kinases p38 and cJun which respectively cause cell cycle arrest and apoptosis (Sinha et al. 2020; Wu et al. 2020).

The idea that ribosome collisions are the essential signal to activate these quality control and stress response pathways is based on four lines of evidence. First, the global induction of ribosome collisions is sufficient to provoke Hel2-dependent ubiquitination of uS3, a hallmark of NGD (Simms et al. 2017b). Second, crucial regulators of quality control responses (Asc1/hRACK1, uS3 and Mbf1/hEDF1) occupy central positions in structures of collided ribosomes (disomes and trisomes), supporting their role in regulation of quality control responses. For instance, yeast Asc1 (hRACK1) (Kuroha et al. 2010) and uS3 (Simms et al. 2018; Wang et al. 2018) reside at the 40S-40S interface of collided ribosomes (Matsuo et al. 2017; Juszkiewicz et al. 2018; Ikeuchi et al. 2019; Sinha et al. 2020), and Mbf1 (hEDF1) (Wang et al. 2018; Simms et al. 2019; Juszkiewicz et al. 2020a; Sinha et al. 2020) interacts with uS3 on the colliding ribosome (Sinha et al. 2020; Pochopien et al. 2021) Third, crucial regulators of quality control are specifically recruited to collided ribosomes, rather than monosomes. For instance, both Hel2 and hZNF598 act preferentially on disomic or trisomic ribosomes *in vitro* (Juszkiewicz et al. 2018; Ikeuchi et al. 2019; Matsuo et al. 2020), and both hZNF598 and hEDF1 are specifically enriched on nuclease-resistant ribosome multimers compared to their relative abundance on monosomes (Juszkiewicz et al. 2020a; Sinha et al. 2020). Gcn1, an essential component of the ISR, also specifically binds collided ribosomes with Mbf1 (Pochopien et al. 2021; Yan and Zaher 2021). Fourth, frameshifting at CGA codon repeats in yeast, which occurs when Mbf1 or uS3 proteins are mutated (Wang et al. 2018), is critically dependent upon ribosome density on the mRNA and the position of the CGA codon repeats relative to the AUG start, as expected if collisions are required for the frameshift (Wolf and Grayhack 2015; Simms et al. 2019).

While the events, components and interactions of the NGD and ISR pathways have been studied extensively, there is far less known about pathways involving Mbf1 (hEDF1) although both proteins affect frameshifting (Hendrick et al. 2001; Wang et al. 2018; Juszkiewicz et al. 2020a). We can infer much about the mechanisms by which Mbf1 prevents frameshifting based upon the structural analyses of collided ribosomes with and without Gcn1 (Sinha et al. 2020; Pochopien et al. 2021). Mbf1 binds the colliding ribosome through interactions with conserved residues of uS3 and also interacts directly with the mRNA entering the colliding ribosome, altering the path of the 3’ end of the mRNA, promoting interactions between the mRNA and h16 of the 40S, and likely locking the 40S head to prevent translocation, all of which are likely to inhibit frameshifting (Sinha et al. 2020; Pochopien et al. 2021). However, as noted above, in mammals frameshifting is much less efficient and occurs in the -1 rather than the +1 direction when ribosomes stall and collide (Juszkiewicz et al. 2020a). Thus, additional factors may promote frameshifting at CGA codon pairs and other inhibitory pairs in yeast (Wang et al. 2018). We note that ribosomes exhibit multiple defects in decoding CGA-CGA and CGA-CCG codons, including a distorted conformation of mRNA in the ribosome A site, both slow and incomplete elongation *in vitro*, and pausing of ribosomes with empty A sites at these pairs *in vivo* (Tesina et al. 2020). Thus, efficient +1 frameshifting at CGA-CGA codon pairs in yeast may be promoted by signals in addition to the ribosome collision, by proteins unique to yeast or by differences between yeast and humans in the relative efficiency of different response pathways, any or all of which could in turn affect the magnitude of frameshifting.

We set out to further understand the forces that promote and inhibit frameshifting and their relationship to other pathways regulated by ribosome collisions. To that end, we selected mutants that suppressed frameshifting when Mbf1 was defective and identified a truncation mutation in the general elongation factor eEF3. We present evidence that the mutant form of eEF3 specifically reduces frameshifting, rather than reducing either inhibition by CGA codon pairs or the overall translation efficiency of an optimal reporter. Thus, we infer that frameshifting is driven by events in addition to the collision, since otherwise effects on stalling and the ensuing collision should equally impact CGA inhibition and frameshifting. We also selected mutants that promoted frameshifting when Mbf1 was functional and found mutations in *GCN1*. Moreover, we uncovered a synergistic interaction between uS3, the site of Mbf1 binding, and Gcn1, which binds collided ribosomes with Mbf1 (Pochopien et al. 2021). We find that Gcn1 modulates frameshifting in conjunction with Gcn20, which also binds collided ribosomes, but that neither Gcn2 nor Gcn4 affect frameshifting, suggesting a new role for Gcn1 and Gcn20 on the collided ribosome distinct from their known role in the ISR pathway. Furthermore, we provide evidence that eEF3 effects on frameshifting are modulated by competition with Gcn1, Mbf1 and the NGD pathway.

## RESULTS

### eEF3 plays a role in frameshifting at CGA codon repeats

Based on the apparent differences between yeast and humans in both directionality and efficiency of frameshifting at collided ribosomes (Wang et al. 2018; Simms et al. 2019; Juszkiewicz et al. 2020a), we considered that frameshifting at CGA codon repeats in yeast might involve yeast-specific factors that promote frameshifting. If so, mutations in the corresponding genes could suppress the frameshifting at CGA codon repeats that is caused by defects in Mbf1. To obtain mutants that suppressed frameshifting caused by *mbf1* mutations, we reversed a previous selection, which yielded the *mbf1* mutants (Wang et al. 2018), using strains in which expression of both the *URA3* and GFP genes require a +1 frameshift downstream of four to six adjacent CGA codons in chromosomally integrated reporters (Fig. 1A). In this background, *mbf1* mutants (Wang et al.2018) exhibited a Ura^+^ GFP^+^ phenotype, since ribosomes frameshift efficiently at CGA repeats in these mutants. To obtain frameshifting suppressors, we selected Ura^-^ mutants based on resistance to 5-fluoro-orotic acid (FOA^R^), since Ura^+^ yeast convert FOA to the toxic compound fluorouracil (Boeke et al. 1984; Boeke et al. 1987). To determine which of the FOA^R^ mutants specifically affected frameshifting, we screened the mutants for reduced expression of the frameshifted GFP reporter, using the ratio of GFP/RFP to account for differences in overall expression of the reporter between strains, as described previously (Dean and Grayhack 2012). To ensure mutants exhibited low levels of frameshifting we identified mutants with GFP/RFP ratios <60% of the parental strain.

**Figure 1.**
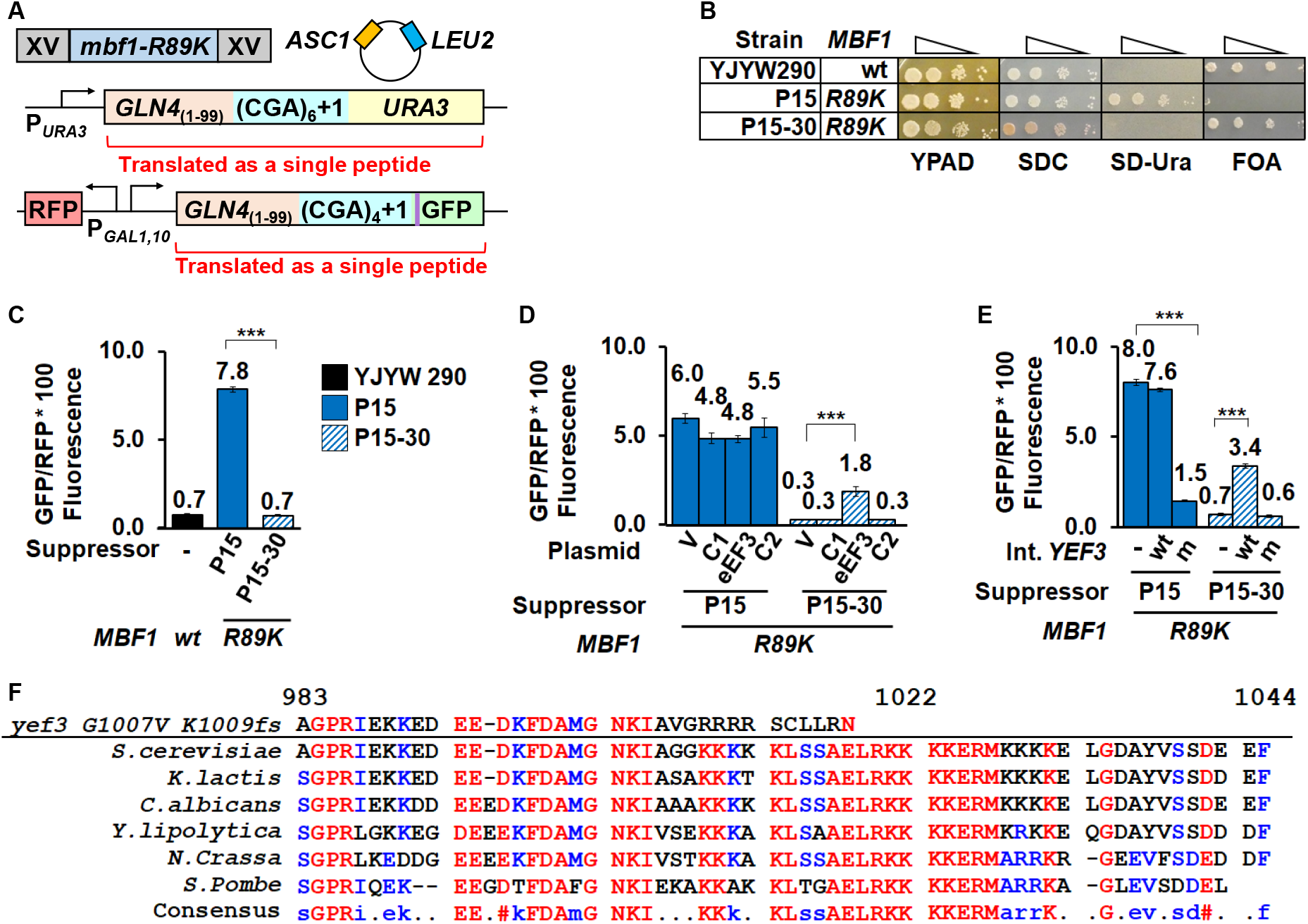
eEF3 plays a role in frameshifting at CGA codon repeats. (A) Schematic of the selection for mutants that suppress frameshifting at CGA codon repeats when Mbf1 is defective. In the P15 selection strain, CGA codon repeats plus a single nucleotide were inserted upstream of the *URA3* and GFP coding regions; the strain also contains a plasmid-borne copy of *ASC1* to avoid mutations in *ASC1.* The *mbf1-R89K* mutant in the P15 strain results in an Ura^+^ GFP^+^ phenotype due to efficient frameshifting at CGA codon repeats (Wang, 2018). Mutants that suppress frameshifting in the *mbf1-R89K* mutant were selected as FOA resistant mutants that also exhibited reduced GFP expression. (B) The P15 suppressor (P15-30) exhibits an Ura^-^ FOA-resistant phenotype, unlike its parent P15, but like its grandparent (YJYW290) [see (Wang et al. 2018)]. Serial dilutions of the indicated strains were grown at 30°C on rich media (YPAD), complete minimal media (SDC), minimal media lacking uracil (SD-Ura), and minimal media containing FOA. (C) Expression of the GLN4_(1-99)_-(CGA)_4_+1-GFP frameshifted reporter is significantly reduced in the P15-30 suppressor relative to its parent (P15). (D) Expression of frameshifted GFP/RFP is partially restored in the P15-30 suppressor by addition of a plasmid bearing the *YEF3* gene (encoding eEF3). GFP/RFP was measured in the P15 parent strain and the P15-30 suppressor strain bearing 2μ plasmids with either no insert (V, vector) or genomic inserts from the yeast tiling collection (Jones et al. 2008). eEF3: plasmid with the *YEF3* gene; C1 and C2: plasmids with flanking chromosomal sequences. (E) Expression of the frameshifted GFP/RFP is modulated by replacement of *YEF3* alleles in the chromosome. In the P15 strain, integration of mutant *yef3-fs1009* into the chromosome results in reduced expression of frameshifted GFP/RFP while in the suppressor P15-30 strain, integration of wild type *YEF3* into the chromosome results in increased expression of frameshifted GFP/RFP. (F) Amino acid sequence alignment of the C-terminal region of eEF3 from six *Ascomycete* fungi across several different clades showing the *yef3 G1007V K1009fs* mutation using MultAlin (http://multalin.toulouse.inra.fr/multalin/) (Corpet 1988). Numbering above the sequences is based on *S. Cerevisiae* eEF3. The color text represents the level of consensus for each residue (Blue: 50-90%, Red: >90%). In all panels, *** indicates p-value of < 0.001.

In this study, we focus on one such mutant, P15-30 which had a mutated version of the gene encoding eEF3 (*YEF3*), based on whole genome sequencing and targeted re-sequencing. In P15-30, the *YEF3* gene bears two mutations, a single amino acid change (G1007V) and a frameshift (K1009fs) which leads to premature termination and loss of 35 amino acids from the C-terminus; we refer to the *yef3-G1007V K1009fs* mutations as *yef3-fs1009* in this paper. As expected if frameshifting is reduced in the P15-30 suppressor, the P15-30 mutant failed to grow on media lacking uracil, unlike its *mbf1^-^* parent P15, but similar to its *MBF1*^+^ grandparent YJYW290 (Wang et al. 2018) (Fig. 1B). Similarly, the P15-30 suppressor and its grandparent exhibited low levels of frameshifted GFP/RFP relative to P15 (Fig. 1C).

To confirm that the *yef3-fs1009* mutation in P15-30 was responsible for the suppression of frameshifting, we first showed that the mutation was recessive, in that a diploid of P15-30 obtained by mating to a *MATα mbf1Δ* strain restored frameshifting to same level as the similarly mated P15 parent (Supplemental Fig. S1). Exogenous expression of wild type *YEF3* (Jones et al. 2008) in the P15-30 suppressor did result in an increase in frameshifted GFP/RFP from 0.3 to 1.8 (Fig. 1D), but did not restore frameshifted GFP/RFP to parental levels of 4.8-5.5 (P15 transformants). To quantitatively determine the effect of the *yef3* mutation on frameshifting suppression, we replaced the chromosomal *yef3-fs1009* allele in P15-30 with wild type *YEF3* kan^R^ and found that expression of the frameshifted reporter increased 5-fold (0.7 to 3.4 GFP/RFP) while replacement with a *yef3-fs1009* kan^R^ construct did not increase GFP/RFP (Fig. 1E). Likewise, we found that replacement of the chromosomal *YEF3* in the P15 parent with *yef3-fs1009* kan^R^ resulted in a 5-fold reduction in GFP/RFP (8.0 to 1.5) (Fig. 1E). Thus, we concluded that the mutation of *YEF3* is both necessary and sufficient to suppress frameshifting in the *mbf1-R89K* mutant.

eEF3 is one of four essential translation factors in yeast, which act during each round of translation to facilitate the steps required for elongation: acceptance of aminoacyl-tRNA into the A site of the ribosome, formation of the peptide bond, translocation of the mRNA with its cognate tRNAs from the A and P sites to the P and E sites, and release of deacyl-tRNA from the E site (Dever and Green 2012; Dever et al. 2016). Unlike the other three elongation factors, which are conserved in all kingdoms, eEF3 is highly conserved (Supplemental Fig. S2) in fungi and other single celled eukaryotes, but has no known homolog in mammals or bacteria (Belfield et al. 1995; Mateyak et al. 2018). eEF3, a member of the ribosome-associated family of ATP-binding cassette (ABC) ATPases (Sandbaken et al. 1990; Andersen et al. 2006; Murina et al. 2019), promotes the late stages of tRNA translocation and facilitates the release of deacyl-tRNA from the E-site (Triana-Alonso et al. 1995; Ranjan et al. 2021). eEF3 is positioned on the ribosome to assist movement of the L1 stalk, providing a structural model for its function of promoting E site release (Andersen et al. 2006; Ranjan et al. 2021). However, the function of the C-terminal domain in which the *G1007V* and *K1009fs* mutations are found is unknown. This domain (residues 981-1044) (Fig. 1F) was not resolved in either structure (Andersen et al. 2006; Ranjan et al. 2021) and is dispensable for the essential function of eEF3 (Anand et al. 2006; Andersen et al. 2006). However, this domain is also reported to have ribosome binding activity (Kambampati and Chakraburtty 1997) and contains three highly conserved lysine blocks (Fig. 1F) that are removed due to the *K1009fs* mutation.

### Translation function and eEF3 levels are affected by the frameshifting suppressor mutant

To determine which aspects of eEF3 function were affected by the *yef3-fs1009* mutation we analyzed the growth of strains with wild type *MBF1* and either wild type *YEF3* or the *yef3-fs1009* mutation. To eliminate the effects of other mutations in the P15 and P15-30 strains, we introduced the *YEF3* and *yef3-fs1009* alleles into wild type BY4741 yeast strains, precisely replacing the chromosomal *YEF3* locus by integrating constructs with *YEF3* (wild type or mutant) fused to a *K. lactis URA5* gene followed by excision of the *K. lactis URA5* marker using selection on FOA-containing media (Boeke et al. 1984; Boeke et al. 1987). We next integrated various *MBF1* alleles into these strains for the experiments described below.

If the *yef3-fs1009* mutation affects an important function of eEF3 in translation, then the *yef3-fs1009* mutation might be expected to alter either growth rate or sensitivity to translation inhibitors. Mutations in *YEF3* (one located between its two ABC domains and one in the chromodomain) are known to result in sensitivity to the aminoglycoside paromomycin (Anand et al. 2003; Sasikumar and Kinzy 2014). Indeed, we found that three independent isolates of strains bearing the *yef3-fs1009* mutation grew slowly on rich media at all temperatures showing an exacerbated growth defect at high temperatures (Fig. 2A). Furthermore, the *yef3-fs1009* mutants were more sensitive at 37°C to both anisomycin, which inhibits peptidyl transferase activity (Grollman 1967) and paromomycin, which relaxes decoding specificity resulting in increased misreading (Fourmy et al. 1996; Fan-Minogue and Bedwell 2008) (Fig. 2B). Thus, we infer that the mutant eEF3 results in a translation defect.

**Figure 2.**
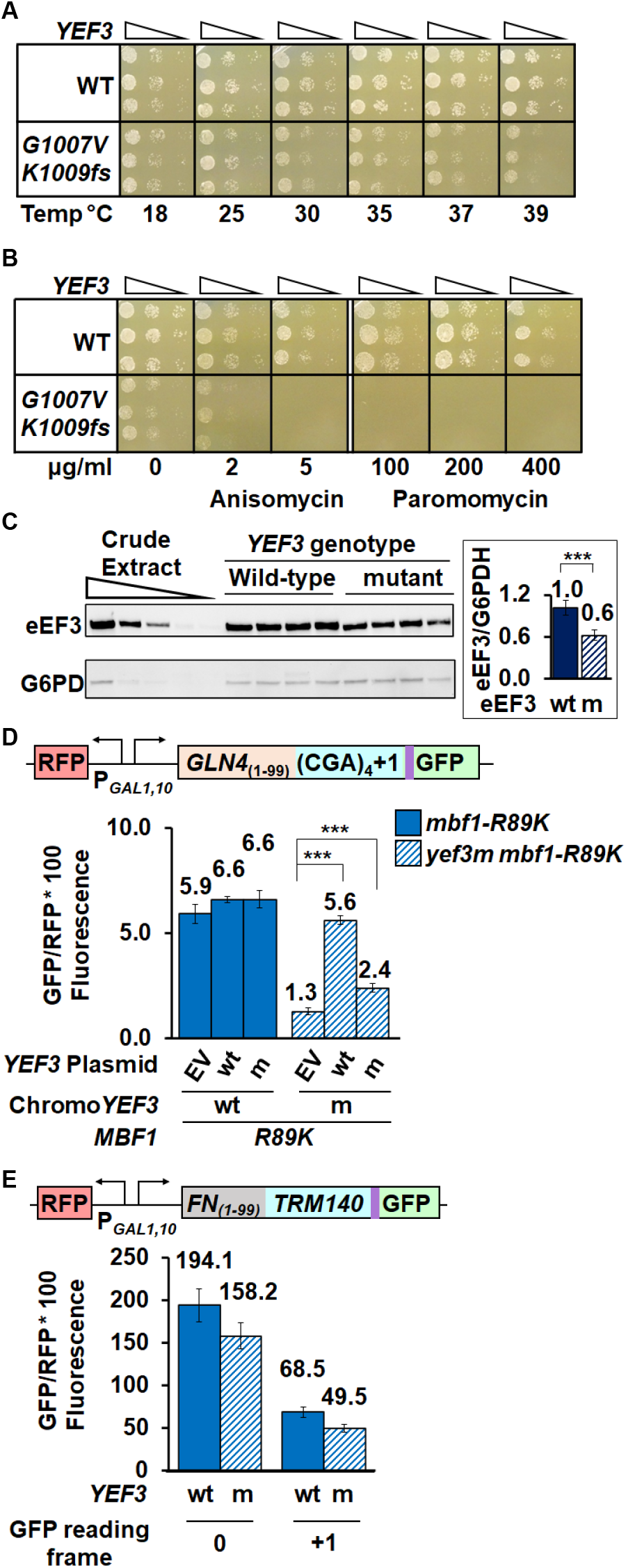
The *yef3-fs1009* mutation results in a temperature dependent growth defect, sensitivity to translation inhibitors and reduced amounts of eEF3. (A) The *yef3-fs1009* mutation results in a growth defect that is exacerbated at higher temperatures. Serial dilutions of *YEF3* wild type and *yef3-fs1009* strains were grown on rich media (YPAD) at the indicated temperatures. (B) The *yef3-fs1009* mutation confers sensitivity to translation inhibitors anisomycin and paromomycin. Serial dilutions of strains in (A) were grown on rich (YPAD) media at 37°C with indicated concentrations of anisomycin or paromomycin. (C) Strains with the *yef3-fs1009* mutation have reduced levels of eEF3 protein compared to otherwise isogenic strains with wild type *YEF3*. Crude extracts, separated by SDS-PAGE, were subjected to Western analysis with anti-eEF3 and anti-glucose-6-phosphate dehydrogenase (G6PDH) antibodies and quantified using Image J (https://imagej.nih.gov/ij/). (D) Increased copies of the *yef3-fs1009* allele result in increased frameshifting in the *yef3-fs1009 mbf1-R89K* mutant, but have no effect in *YEF3 mbf1-R89K* strains. GFP/RFP expression from the (CGA)_4_+1 reporter was examined in *YEF3* and *yef3-fs1009* strains bearing *CEN* plasmids with no insert (EV), YEF3 (wt) or *yef3-fs1009* (m). (E) The *yef3-fs1009* mutation does not suppress programmed frameshifting in *TRM140* mRNA. ** indicates a p-value <0.01 ≥ 0.001 *** indicates p-value of < 0.001.

Since the C-terminal domain of eEF3 itself is not essential (Andersen et al. 2006), we considered that mutant phenotypes might be caused by reduced eEF3 protein levels. Indeed, we found a reduction in both antibody-reactive eEF3 in the mutant, to 60% of that in wild type (Fig. 2C) and an apparent reduction in size and intensity of a Coomassie-stained band likely corresponding to eEF3 (Supplemental Fig. S3A).

To determine if the frameshifting suppression observed in the *yef3-fs1009* mutant was due to limiting amounts of eEF3 protein, we examined the effect of increased expression of the mutant and wild type eEF3 on frameshifted GFP/RFP and on eEF3 levels. As expected, expression of wild type *YEF3* from a *LEU2 CEN* plasmid in the *yef3-fs1009* mutant nearly completely restored expression of the frameshifted reporter from 1.3 GFP/RFP with the empty vector to 5.6 GFP/RFP (a 4.3-fold increase), 95% of that in a *YEF3* wild type strain with an empty vector (Fig. 2D). Thus, the mutant is completely recessive. By contrast, increased expression of *yef3-fs1009* to levels exceeding those in wild type (Supplemental Fig. S3B) had a much reduced effect, resulting in 2.4 GFP/RFP (a 1.8-fold increase relative to the vector control), 41% of that in a *YEF3* wild type strain with an empty vector (Fig. 2D). We conclude that the effects of the *yef3-fs1009* mutant are due to two effects, one that is due to reduced expression and a separate effect that is due to reduced function (and cannot be restored with increased amounts).

Since the E site at which eEF3 acts (Triana-Alonso et al. 1995; Ranjan et al. 2021) has been implicated in programmed frameshifting (Marquez et al. 2004; Sanders and Curran 2007; Devaraj et al. 2009), we considered that the *yef3-fs1009* mutant might primarily affect frameshifting. To test this idea, we compared the frameshifting efficiency of two native yeast +1 frameshifting signals, *TRM140* and *TY1*, in wild type yeast and the *yef3-fs1009* mutant. While the site of frameshifting at these two sites is identical, CUU-AGG-C (Belcourt and Farabaugh 1990; Asakura et al. 1998; Farabaugh et al. 2006), the *TRM140* site is a remarkably efficient frameshifting signal (D’Silva et al. 2011), perhaps due to sites upstream of the frameshift at which ribosomes collide (Meydan and Guydosh 2020). We observed highly efficient *TRM140* frameshifting in both wild type and *yef3-fs1009* mutant strains based on frameshifted GFP/RFP levels relative to in-frame expression, 68.5 compared to 194.1 in wild type (35%) and 49.5 compared to 158.2 in the mutant (31%) (Fig. 2E). Thus, there was little difference in relative frameshifting between *YEF3* wt and the *yef3-fs1009* mutant. Similarly, there was little difference in frameshifted GFP/RFP with the *TY1* signal although the level of frameshifted protein was much less than with *TRM140* (Supplemental Fig. S3C). Thus, suppression of frameshifting at CGA codon repeats is unlikely due to an inherent defect in frameshifting ability.

### eEF3 modulates frameshifting caused by defects in either Mbf1 or ribosomal protein S3

Reading frame maintenance at collided ribosomes depends upon the extra-ribosomal protein Mbf1 and its interaction with uS3 in the colliding ribosome, as mutations in *RPS3* (encoding yeast uS3) that affect this interface result in frameshifting (Wang et al. 2018; Juszkiewicz et al. 2020a; Sinha et al. 2020; Pochopien et al. 2021). To ascertain the nature of the suppression by the *yef3-fs1009* mutant, we examined both the types of frameshifting mutations suppressed by *yef3-fs1009* as well as the effects of *yef3-fs1009* on the levels of frameshifted protein (GFP fluorescence), mRNA and the ratio of GFP to mRNA from the *GLN4*_(1- 99)_-(CGA)_4_+1-GFP reporter (Fig. 3A). To this end, we examined the ability of the *yef3-fs1009* mutation to suppress frameshifting in four mutants: *mbf1-R89K*, *mbf1Δ, RPS3-K108N* and *RPS3-S104Y*.

**Figure 3.**
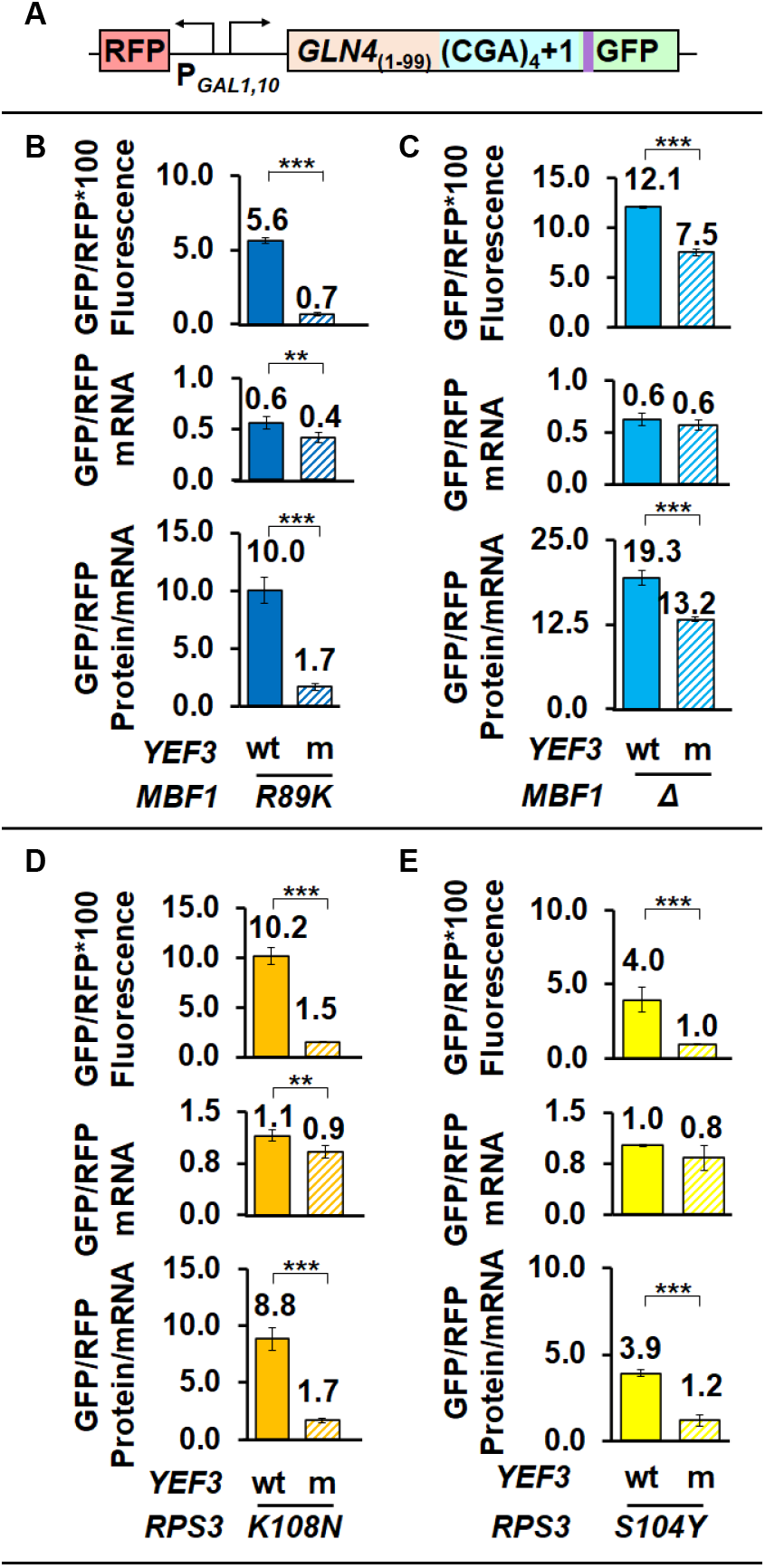
The *yef3-fs1009* mutation suppresses frameshifting at CGA codon repeats when the frame quality control system is compromised by defects in *MBF1* or *RPS3*. (A) Schematic of RFP and *GLN4*_(1-99)_-(CGA)_4_+1-GFP reporter used in these analyses. (B, C, D, E) The *yef3-fs1009* mutation suppresses frameshifting caused by the *mbf1-R89K* mutation (B), by deletion of *MBF1* (C), by the *RPS3-K108N* mutation (D) or by the *RPS3-S104Y* mutation (E), In each case, the *yef3-fs1009* mutation results in significantly reduced levels of both frameshifted protein and protein per mRNA and in two cases results in small but significant reductions in mRNA. ** indicates a p-value <0.01 ≥ 0.001 *** indicates p-value of < 0.001.

The *yef3-fs1009* mutant suppressed frameshifting caused by mutations in either *MBF1* or *RPS3*, albeit with some differences in effectiveness. The *yef3-fs1009* mutant had major effects on frameshifted protein per mRNA for each of the three point mutations (*mbf1-R89K*, *RPS3-K108N* and *RPS3-S104Y*). In the *mbf1-R89K* strains, the *yef3-fs1009* mutation resulted in a 5.9-fold reduction in frameshifted GFP/RFP per mRNA (10.0 to 1.7) (Fig. 3B). Similarly, in the *RPS3-K108N* and *RPS3-S104Y* strains, the *yef3-fs1009* mutation resulted in 5.2 and 3.3-fold reductions in frameshifted GFP/RFP per mRNA (8.8 to 1.7 and 3.9 to 1.2) (Fig. 3D and 3E). By contrast, in *mbf1Δ* strains, the *yef3-fs1009* mutation had a much smaller 1.5-fold effect on frameshifted GFP/RFP (19.3 in the *YEF3 mbf1Δ* strain compared to 13.2 in the *yef3-fs1009 mbf1Δ* strain). We noted that in most cases, the *yef3* mutant also exhibited a small (but in some cases significant) reduction in mRNA levels (Fig. 3 B-E), which could indicate increased mRNA decay. However, in no case did the reduction in mRNA account for the reduced amount of frameshifted protein, thus we conclude that *yef3-fs1009* suppressed ribosomal frameshifting at CGA codon repeats in a manner independent of the gene or the particular mutation that allowed that frameshifting. The reduced effectiveness of the *yef3* mutant in the complete absence of Mbf1 protein is consistent with the idea the Mbf1 and eEF3 compete for the collided ribosome (see below).

### eEF3 has specific effects on frameshifting, rather than CGA inhibition

eEF3 acts in each cycle of translation and its depletion in yeast cells altered both the rate-limiting step in translation and codon discrimination (Ranjan et al. 2021). Thus, we considered that the *yef3-fs1009* mutant could exert its effects by altering the stall at CGA-CGA codon pairs, or by altering overall ribosome availability and thus impacting the frequency of the ribosome collisions that lead to both inhibition and frameshifting (Simms et al. 2017a; Simms et al. 2019). To test these possibilities, we examined the effects of the *yef3-fs1009* mutant on in-frame expression of reporters with inhibitory (CGA-CGA) codon pairs and the corresponding optimal (AGA-AGA) codon pairs. We expected to observe a reduction in CGA inhibition in the mutant strain, if either the stall or collisions at CGA-CGA codon pairs were reduced in the *yef3-fs1009* mutant. We might observe a reduction in expression of the optimal reporter, if the overall rate of initiation was reduced in the *yef3-fs1009* mutant. We performed these experiments in reconstructed strains bearing *mbf1-R89K* mutations to assess frameshifting of a related reporter in parallel.

We found little difference between the *yef3-fs1009* and wild type *YEF3* strains in the expression of either the inhibitory or optimal in-frame reporters (Fig. 4A). CGA inhibition as measured by GFP levels from the CGA reporter relative to those from the AGA reporter were 35% in the*YEF3* wild type strain (23.8 to 68.1 GFP/RFP fluorescence) and 32% in the *yef3-fs1009* strain (21.9 to 69.3 GFP/RFP fluorescence) (Fig. 4A); similarly, GFP/RFP protein per mRNA levels were 43% (25.6 to 59.7) and 46% (20.2 to 44.1) respectively. As expected, the *yef3-fs1009* mutation suppressed frameshifting in the *GLN4*_(1-99)_-(CGA-CGA)_3_ +1-GFP reporter (6.9 to 3.1 GFP/RFP protein per mRNA) (Fig. 4A). We verified that CGA inhibition was also substantial in *yef3-fs1009* strains bearing either wild type MBF1 (Supplemental Fig. S4A) or *mbf1Δ* (Supplemental Fig. S4B). Thus, CGA-CGA codon pairs are inhibitory in the *yef3-fs1009* strains, suggesting that ribosomes stall and collide in the mutant strain, consistent with a specific defect in frameshifting caused by the mutation.

**Figure 4.**
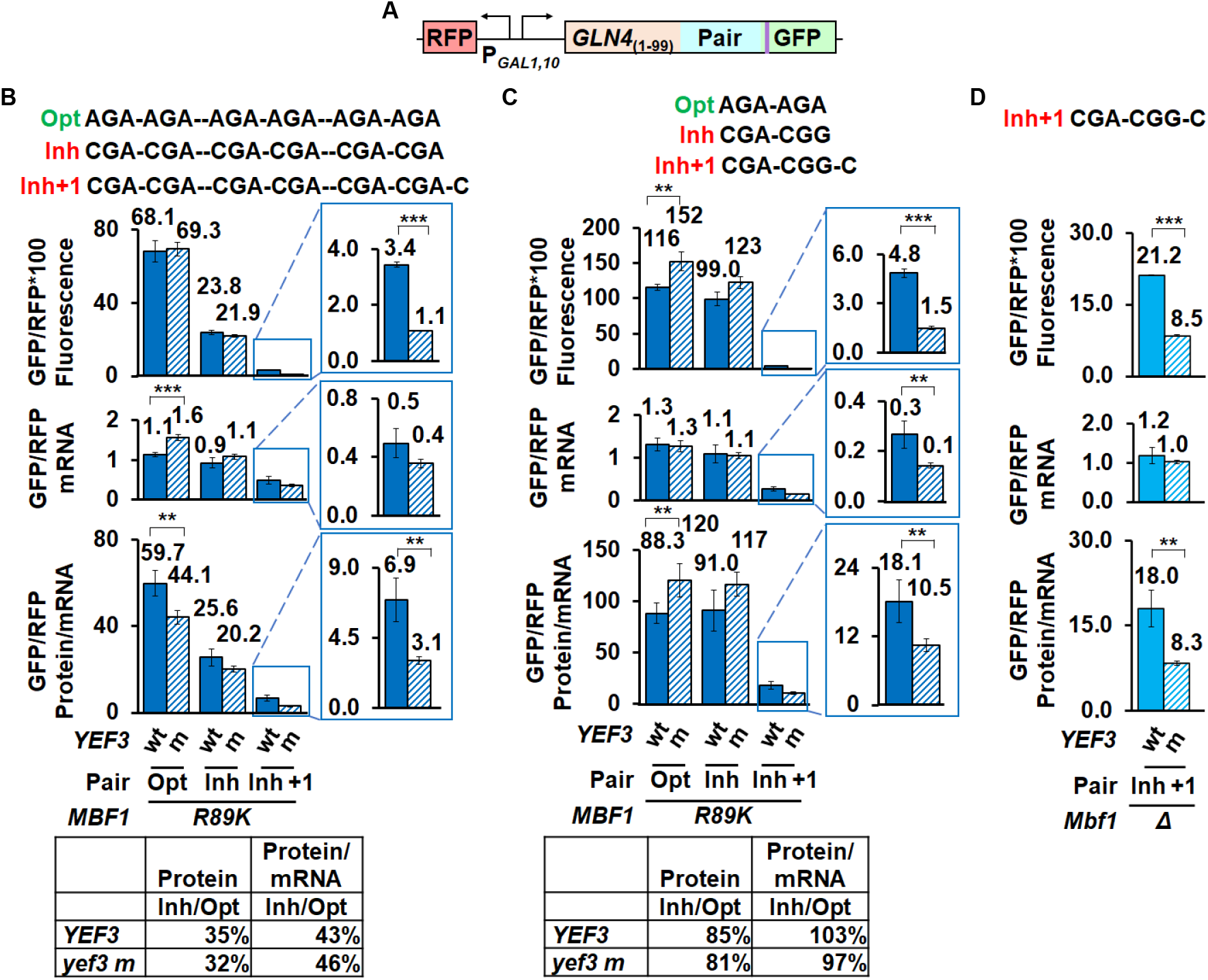
The *yef3-fs1009* mutation suppresses frameshifting with two different inhibitory codon combinations, but has only small effects on in-frame expression of reporters with optimal or inhibitory codons. (A) The *yef3-fs1009* mutation suppresses frameshifting in *mbf1-R89K* mutant bearing a reporter with three copies of CGA-CGA codon pairs but does not relieve or enhance CGA inhibition from in-frame reporters. Levels of GFP/RFP protein (fluorescence), mRNA and protein/mRNA are similar from in-frame reporters with optimal (AGA-AGA) or inhibitory (CGA-CGA) codon pairs, but levels of protein and protein/mRNA levels are significantly different from the analogous frameshifting reporter (CGA-CGA)_3_+1 insert. In-frame expression of inhibitory (CGA) reporter relative to optimal (AGA) reporter is shown below. (B) The *yef3-fs1009* mutation suppresses frameshifting in an *mbf1-R89K* mutant with a reporter bearing a single CGA-CGG inhibitory pair, but does not affect CGA-CGG inhibition. In-frame expression of inhibitory (CGA-CGG) reporters relative to optimal (AGA-AGA) reporters is similar in *YEF3* and the *yef3-fs1009* mutant strains (table), although levels of GFP/RFP protein and protein/mRNA from both in-frame reporters are greater in the *yef3-fs1009* mutant. By contrast, both GFP/RFP protein and protein/mRNA levels from the frameshifting reporter are reduced in the *yef3-fs1009* mutant. (C) The *yef3-fs1009* mutation suppresses frameshifting in an *mbf1Δ* mutant with a reporter bearing a single CGA-CGG inhibitory pair. ** indicates a p-value <0.01 ≥ 0.001 *** indicates p-value of < 0.001.

One might consider that the primary effect of the mutant eEF3 protein at CGA-CGA codon pairs is to allow ribosomes to abort translation rather than frameshift, since a large fraction of ribosomes fail to translate through the strongly inhibitory sequences (Matsuo et al. 2017; Sitron et al. 2017). To assess whether the mutant eEF3 really suppresses frameshifting or simply results in stalled ribosomes aborting translation, we examined the ability of the *yef3-fs1009* mutant to suppress frameshifting at a high frameshifting site with minimal inhibitory effects. We had previously determined that ribosomes frameshift efficiently at a single CGA-CGG-C site (Wang et al. 2018). We found that this CGA-CGG-C site is minimally inhibitory, in that expression of the in-frame GFP/RFP protein with the inhibitory codon pair is 85% that of the reporter with the optimal codon pair in the wild type strain (99.0 to 116 GFP/RFP fluorescence) and 81% in the *yef3-fs1009* mutant (123 to 152 GFP/RFP fluorescence) (Fig. 4B) [compared to 35% (23.8 to 68.1) and 32% (21.9 to 69.3) respectively with the (CGA-CGA)_3_ reporter (Fig. 4A)]. Nevertheless, we find that the frameshifted GFP/RFP protein at the CGA-CGG-C site is significantly reduced in the *yef3-fs1009* mutant (1.5 GFP/RFP fluorescence) relative to the *YEF3* wt (4.8 GFP/RFP fluorescence) (Fig. 4B). Suppression is still apparent in the frameshifted protein/mRNA, although it is clear that significant reduction in mRNA also occurred in the *yef3-fs1009* mutant relative to the wild type (Fig. 4B). We also examined frameshifting suppression at this site in a set of *mbf1Δ* mutants, since the original suppression in these mutants had been less robust. Again, we found that both frameshifted protein and protein per mRNA were significantly reduced in the *yef3-fs1009* mutant (Fig. 4C). Thus, we infer that frameshifting suppression caused by this mutant eEF3 is likely not dependent upon aborting translation at a high rate.

### Integrated Stress Response Regulators Gcn1 and Gcn20 inhibit frameshifting at CGA codon repeats

To understand frameshifting during ribosome collisions, we previously obtained mutants that allow frameshifting at CGA codon repeats in wild type strains and identified mutations in *MBF1* and *RPS3* (Wang et al. 2018). To identify additional genes in this process, we repeated the selection and screen for Ura^+^ GFP^+^ mutants in an *MBF1* strain in which expression of both *URA3* and GFP require frameshifting (Supplemental Fig. S5A) (Wang et al. 2018), but which also carried plasmid-borne copies of both *MBF1* and *ASC1* to avoid recessive mutations in these genes. Among the Ura^+^ GFP^+^ mutants that passed this screen, whole genome sequencing yielded seven independent mutations in *GCN1* (Supplemental Table S1). *GCN1* encodes a key regulator of the Integrated Stress Response pathway (Garcia-Barrio et al. 2000; Sattlegger and Hinnebusch 2000; Hinnebusch 2005), known to bind to polyribosomes (with Gcn20) as an essential component of its activation of Gcn2, the eIF2α kinase that initiates the ISR (Marton et al. 1997; Sattlegger and Hinnebusch 2005). The role of Gcn1 in reading frame maintenance at collided ribosomes is particularly interesting for two reasons: 1. Gcn1 and Gcn20 bind collided ribosomes with Mbf1 (Pochopien et al. 2021), and 2. Gcn1 and Gcn20 share extensive homology with eEF3 in their ribosome binding domains and appear to compete with each other for binding to the ribosome (Marton et al. 1993; Visweswaraiah et al. 2012).

To study the role of Gcn1 in frameshifting at CGA codon repeats, we constructed *gcn1Δ* mutants and assayed frameshifting with our (CGA)_4_+1 frameshifting reporter (Fig. 5 A). We observed an increase in frameshifted GFP/RFP in the *gcn1Δ* strains (from 0.5 to 1.9 GFP/RFP), but serendipitously discovered an amplification of frameshifting in *RPS3-S104Y gcn1Δ* mutants, a 6-fold increase over that seen with the *RPS3-S104Y* mutation alone (from 3.3 to 20.2 GFP/RFP) (Fig. 5B). The expression of *GCN1* on a *LEU2 CEN* plasmid in the *gcn1Δ* or *gcn1Δ RPS3-S104Y* strains returned GFP/RFP expression of the frameshifted reporter to levels observed in the wild type or *RPS3-S104Y* strains, but had no effect on the GFP/RFP in either wild type or *RPS3-S104Y* strains (Fig. 5B). Similarly, overexpression of wild type *RPS3* partially suppressed frameshifting in strains with the *RPS3-S104Y* mutation, while expression of the *RPS3-K108E* mutated uS3 exacerbated frameshifting in all strains, particularly those with a *gcn1Δ* mutation (Supplemental Fig. S5B), providing additional evidence that the mutated uS3 protein is responsible for the enhanced frameshifting. We demonstrated that the increase in frameshifted protein was indeed due to an increase in frameshifting rates rather than stabilization of the mRNA, since levels of frameshifted protein per mRNA increased from 2.2 and 7.3 in the *gcn1Δ* and *RPS3-S104Y* single mutants to 25.7 in the double mutant (Fig. 5C). We confirmed that Gcn1 and uS3 proteins have specific roles of in frameshifting, by showing, as we did above for the *yef3-fs1009* mutants, that the *gcn1Δ* and *RPS3-S104Y* mutants have little effect on CGA inhibition with in-frame reporters (Supplemental Fig. S5C).

**Figure 5.**
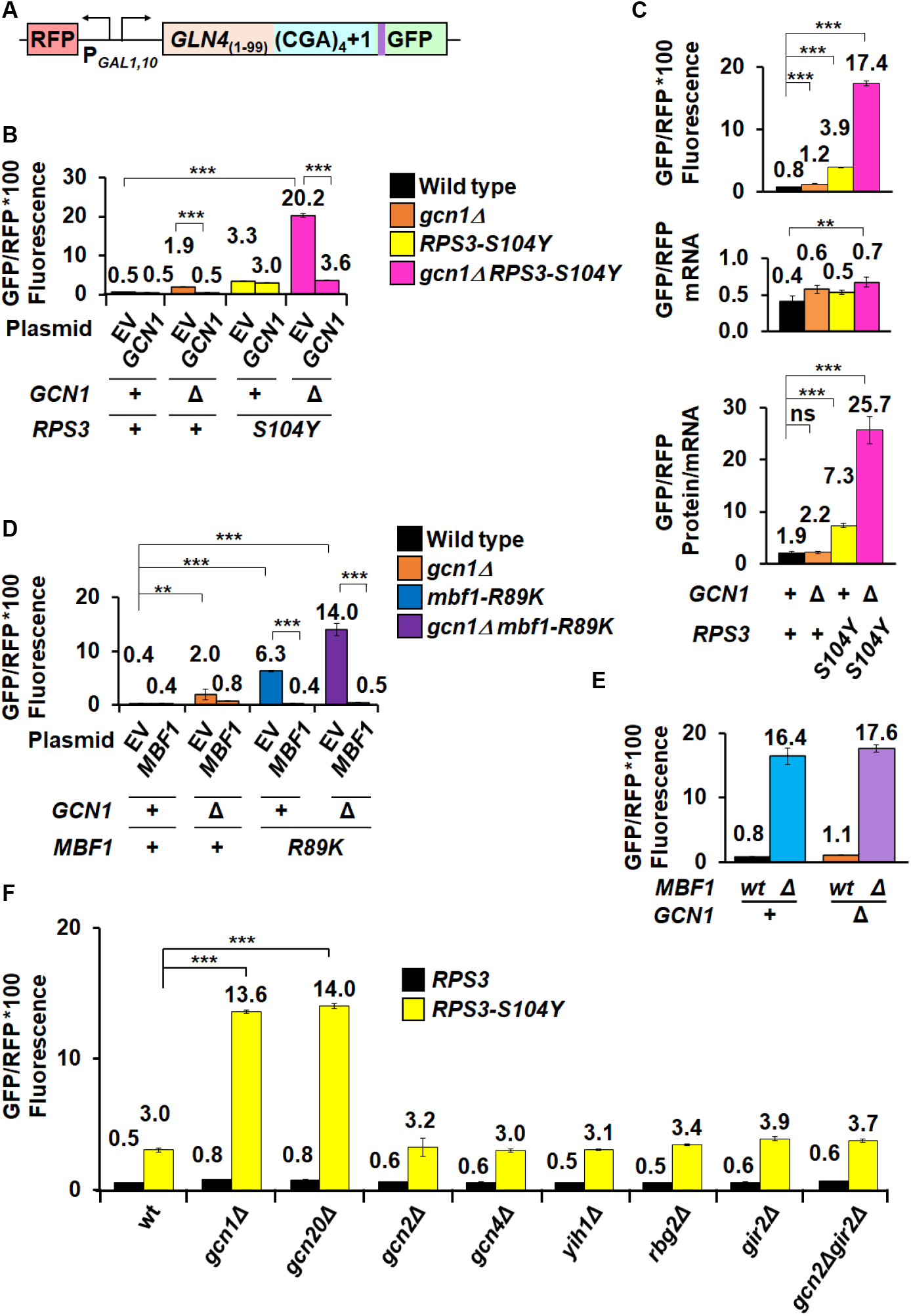
Gcn1 and Gcn20 antagonize frameshifting, but their effects are not mediated by the ISR pathway. (A) Schematic of RFP and *GLN4*_(1-99)_-(CGA)_4_+1-GFP reporter used in these analyses. (B) Deletion of *GCN1* combined with the *RPS3-104Y* mutation results in increased expression of the frameshifted reporter relative to either single mutation. Expression of *GCN1* suppresses the frameshifting in the *gcn1Δ* mutant and in the *gcn1Δ RPS3-S104Y* double mutant but has no effect on frameshifting in the *RPS3-S104Y* single mutant. (C) The increase in frameshifting in the *gcn1Δ RPS3 S104Y* double mutant is not due to effects on mRNA levels. The *gcn1Δ RPS3 S104Y* double mutant significantly increases expression of both GFP/RFP protein and protein per mRNA but has only small effects on the mRNA for the frameshifted reporter. (D) Deletion of *GCN1* combined with the *mbf1-R89K* mutation results in increased expression of the frameshifted reporter relative to either single mutation. Expression of *MBF1* suppresses frameshifting in both the *mbf1-R89K* mutant and in the *gcn1Δ mbf1-R89K* double mutant. (E) Deletion of *GCN1* has only minor effects on the expression of the frameshifted reporter in combination with a deletion of *MBF1*. (F) Deletion of either *GCN1* or *GCN20* in an RPS3-S104Y mutant results in increased expression of the frameshifted reporter. However, deletions ofgenes encoding other components of the ISR pathway (*GCN2, GCN4, YIH1*) or proteins that modulate the ISR pathway and interact with Gcn1 on the collided ribosome (*GIR2, RBG2*) have minimal effects on frameshifting. ** indicates a p-value <0.01 ≥ 0.001 *** indicates p-value of < 0.001.

Similarly, we observed increased frameshifting with *gcn1Δ mbf1-R89K* double mutants (14.0 GFP/RFP), compared to either single mutant (2.0; 6.3 GFP/RFP), which was similarly complemented by expression of *MBF1* (Fig. 5D). By contrast, we did not observe increased frameshifting in *gcn1Δ mbf1Δ* mutants relative to *mbf1Δ* mutants alone (Fig. 5E). Thus, Mbf1 protein must be present for Gcn1 to modulate frameshifting.

Since Gcn1 is a key component of the ISR pathway, we examined the effects of other components in this pathway to find out if frameshifting is modulated by induction of the pathway. Induction of the ISR pathway involves Gcn1 and Gcn20-dependent activation of the Gcn2 kinase, which in turn phosphorylates eIF2α, reducing translation initiation and causing induction of the Gcn4 transcriptional regulator which modifies expression of more than 500 yeast genes (Jia et al. 2000; Natarajan et al. 2001). Moreover, two other proteins interact with Gcn1 on collided ribosomes: Gir2, a competitor of Gcn2, and Rbg2, a ribosome binding GTPase, (Wout et al. 2009; Pochopien et al. 2021). If frameshifting depends upon the induction of the ISR pathway, we expected that deletion of some of these components would yield a synergistic increase in frameshifted GFP/RFP in an *RPS3-S104Y* mutant. We measured frameshifted GFP/RFP in both wild type and *RPS3-S104Y* mutants with deletions in various ISR components (Fig. 5F). Only deletions of *GCN1* or *GCN20* resulted in significantly increased expression of the frameshifted reporter when combined with *RPS3-S104Y,* yielding 13.6 GFP/RFP and 14.0 GFP/RFP respectively compared to 3.0 GFP/RFP in the *RPS3-S104Y* mutant strain (Fig. 5F). While deletion of *GCN20* resulted in high levels of frameshifting, which were complemented by a plasmid-borne copy of *GCN20* (Supplemental Fig. S5D), deletion of the other two genes encoding proteins (Gir2 or Rbg2) that bind the leading stalled ribosome had only small effects on frameshifting (Fig. 5F). We also examined effects of a combined deletion of *GIR2* and *GCN2* since these two proteins are thought to compete for ribosome access (Wout et al. 2009), but their combined deletion had no greater effect on the expression of the frameshifted reporter than the *gir2Δ* mutant (Fig. 5F). Furthermore, we tested the possibility that *gcn1Δ* exerted its effects by allowing spurious activity of the Gcn2 kinase, but found that *gcn1Δ gcn2Δ RPS3-S104Y* triple mutants exhibited the high frameshifting seen in *gcn1Δ RPS3-S104Y* mutants (Supplemental Fig. S5E). Thus, we conclude that Gcn1 and Gcn20 exert their effects on frameshifting through the complex on the ribosome and not through the ISR pathway or through the Gir2 and Rbg2 proteins that bind with collided ribosomes with them.

### eEF3 and Quality Control pathways compete to regulates frameshifting

We considered a model in which eEF3 competes with Hel2 and the Gcn1/Gcn20 complex to act on the collided ribosome. The idea stems from two observations: 1. the NGD and ISR pathways are in competition with each other, such that an increase in activation of the ISR occurs if Hel2 is missing (Meydan and Guydosh 2020; Yan and Zaher 2021) and 2. eEF3 and Gcn1/20 compete in activation of the ISR, since overproduction of eEF3 reduces ISR activation (Visweswaraiah et al. 2012). Specifically, we thought that eEF3, Hel2 and Gcn1 might each compete for collided ribosomes, determining the stalled ribosome’s fate, and the mutant eEF3 protein might not compete well. If so, we expected that efficient frameshifting would be restored in the *yef3-fs1009* mutant if the appropriate pathway(s) was inactivated. To this end, we constructed *hel2Δ* and *gcn1Δ* single mutants and *hel2Δ gcn1Δ* double mutants in *YEF3* and *yef3-fs1009* strains and assessed frameshifting in these strains (Fig. 6A and 6B). Indeed, we found that levels of frameshifted protein in the *hel2Δ gcn1Δ yef3-fs1009* mutant were both high (19.6 GFP/RFP) and fairly similar to that in the corresponding *hel2Δ gcn1Δ YEF3* strain (24.5 GFP/RFP) (80%) (Fig. 6B). By contrast, in the *HEL2^+^ GCN1^+^* parent *yef3-fs1009* mutant, frameshifting was low (1.8 GFP/RFP) and only 26% that of its corresponding *YEF3* strain (1.8 to 6.8 GFP/RFP). Thus, ribosomes using this mutant eEF3 can frameshift if competing pathways are blocked.

**Figure 6.**
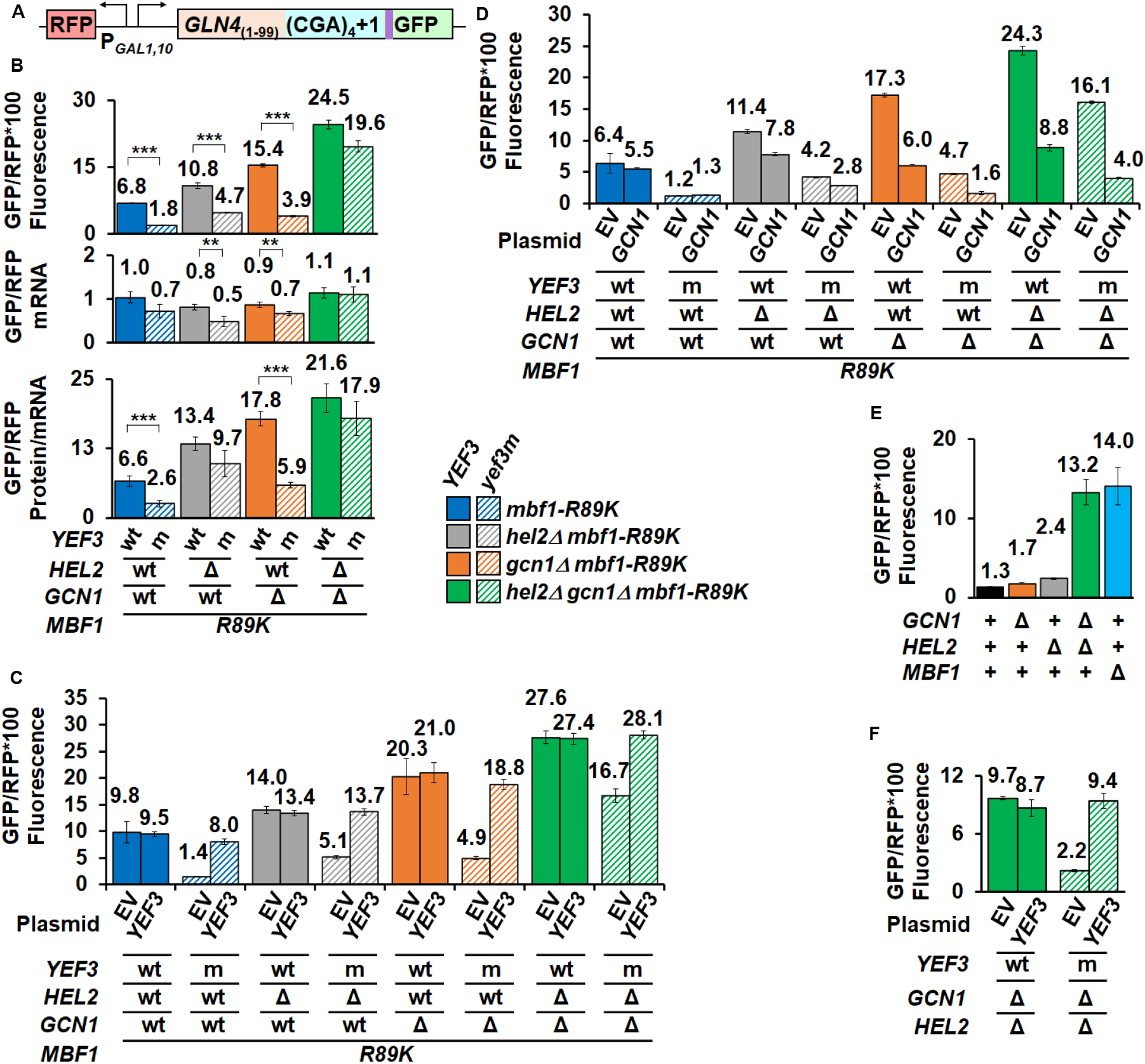
eEF3 competes with Hel2 and Gcn1 to regulate frameshifting at CGA codon repeats. (A) Schematic of RFP and *GLN4*_(1-99)_-(CGA)_4_+1-GFP reporter used in these analyses (B) The suppression of frameshifting due to the *yef3-fs1009* mutation is nearly lost in mutants lacking both *HEL2* and *GCN1*. Similar effects are observed with removal of *HEL2* alone. (C,D) Complementation of mutants with plasmid-born *YEF3* (C) or *GCN1* (D). (E) High levels of frameshifting are observed in *hel2Δ gcn1Δ* mutants despite the presence of a functional *MBF1* gene. (F) The *yef3-fs1009* mutation suppresses the high levels of frameshifting in the *gcn1Δ hel2Δ* mutant. Frameshifting suppression in the *yef3-fs1009* mutant is complemented with addition of a *CEN* plasmid bearing *YEF3*.

We also note that Hel2 appears to play a larger role in the competition than Gcn1. The deletion of *HEL2* resulted in an increase in the relative frameshifted protein in the *hel2Δ yef3-fs1009* mutant to 43% that of the corresponding *YEF3* strain (4.7 to 10.8 GFP/RFP) (Fig. 6B). Furthermore, the *hel2Δ* mutants also displayed differences in mRNA levels that further increased the apparent frameshifting per mRNA in the *hel2Δ yef3-fs1009* mutant (9.7 to 13.4 GFP/RFP/mRNA) (Fig. 6B). By contrast, deletion of *GCN1* resulted in no increase in the relative frameshifted protein (26%; 3.9 to 15.4 GFP/RFP), but these interpretations are complicated because the *gcn1Δ* mutants also display significantly higher levels of frameshifting than the parents.

In introducing these multiple mutations, we were concerned that strains acquired secondary mutations that are responsible for some of these phenotypes. Therefore, we complemented each of the mutations in these strains (Fig 6D and 6E; Supplemental Fig. S6A). As expected, expression of wild type *YEF3* in all strains with *yef3-fs1009* mutation resulted in increased frameshifting to similar levels as the corresponding *YEF3* strain (Fig. 6C). Expression of either *GCN1* or *HEL2* resulted in reduced frameshifting in the corresponding *gcn1Δ* or *hel2Δ* mutants (Fig. 6D; Supplemental Fig. S6A). Surprisingly, expression of *GCN1* also reduced frameshifting in *hel2Δ* mutants (from 11.4 to 7.8 GFP/RFP), but not in wild type (Fig. 6D); likewise, expression of *HEL2* reduced frameshifting in *gcn1Δ* mutants (from 17.4 to 13.4 GFP/RFP) (Supplemental Fig. S6A). The apparent cross complementation is consistent with a strong competition between the pathways, as suggested by previous results (Meydan and Guydosh 2020; Yan and Zaher 2021). Perhaps most surprisingly, we found that while expression of wild type *MBF1* fully suppressed frameshifting in *HEL2 GCN1* or single mutant strains, expression of *MBF1* only partially suppressed frameshifting in *gcn1Δ hel2Δ* mutants bearing either *YEF3* wild type or the *yef3-fs1009* mutation (Supplemental Fig. S6B).

Given the poor suppression of frameshifted protein by *MBF1* wild type in the *gcn1Δ hel2Δ mbf1-R89K* mutants (Supplemental Fig. S6B), we considered the possibility that Mbf1 is unable to prevent frameshifting when neither Hel2 nor Gcn1 is present. We tested this idea and indeed found very high levels of frameshifting in a *gcn1Δ hel2Δ MBF1* strain (Fig. 6E). Thus, Mbf1 presumably requires either the Gcn1/Gcn20 complex or Hel2 to prevent frameshifting.

To find out if eEF3 competes with Mbf1 when the Gcn1/Gcn20 and Hel2 branches are blocked, we asked if frameshifting in a *gcn1Δ hel2Δ MBF1* strain was modulated by *YEF3*. Indeed, this *yef3-fs1009* mutant strain exhibited reduced levels of frameshifted GFP/RFP (2/2 compared to 9.7), which were restored by expression of wild type *YEF3* (Fig. 6F). Overall, we infer that eEF3, Hel2, Gcn1, and Mbf1 compete for collided ribosomes, with each regulator setting off distinct events, and that Mbf1 relies on Hel2 or Gcn1 to act on and remove ribosomes that would otherwise frameshift. Moreover, eEF3 is integral to the frameshifting event.

## DISCUSSION

We showed here that frameshifting at collided ribosomes is provoked by the general translation elongation factor eEF3 and is restrained by multiple aspects of the quality control systems, including not only Mbf1, but also the ISR regulators Gcn1 and Gcn20, and the NGD regulator Hel2. We deduce that wild type eEF3 protein is required for frameshifting at CGA codon repeats, based primarily on the finding that the *yef3-*fs1009 mutation in the gene encoding eEF3 suppresses frameshifting at CGA codon repeats when Mbf1 is defective. We infer that eEF3 has a specific role in frameshifting, rather than simply mediating its effects on frameshifting through effects on ribosome stalls or collisions, based on two observations. First, the *yef3-fs1009* mutant does not affect CGA-CGA inhibition, an argument that the *yef3-fs1009* mutant specifically affects frameshifting, rather than the ribosome collisions or stalls that are necessary for both frameshifting and CGA-CGA inhibition (Letzring et al. 2013; Simms et al. 2017b; Sitron et al. 2017; Simms et al. 2019). Second, the *yef3-fs1009* mutant also suppresses frameshifting at a site that produced little overall inhibition of expression, an argument that neither aborted translation nor strong inhibition are required for the *yef3-fs1009* mutant’s effects. We infer that eEF3 effects do not depend upon any specific quality control component that inhibits frameshifting, as the *yef3-fs1009* mutant suppressed frameshifting in mutants that were simultaneously defective in two of the three quality control regulators (*hel2Δ gcn1Δ* and *mbf1-R89K gcn1Δ* double mutants), Finally, we argue that the *yef3-fs1009* mutant’s effects are not due to a specific interaction with the Mbf1-R89K protein, since the *yef3-fs1009* mutant suppressed frameshifting caused by different defects in either Mbf1 or uS3. Thus, the general elongation factor eEF3 is specifically required for frameshifting at collided ribosomes. Our results provide the first evidence of a direct involvement of the general translation apparatus in the events occurring when ribosomes collide, and the first evidence of a unique role for eEF3 in ribosome collisions.

The involvement of eEF3 in frameshifting in yeast may help to explain differences between yeast and humans in the magnitude and directionality of frameshifting during ribosome collisions (Juszkiewicz et al. 2020a), since mammals do not have an eEF3 homolog (Belfield et al. 1995; Mateyak et al. 2018). For example, an eEF3 compensatory mechanism in mammals may not actively participate in frameshifting.

The relationships between different pathways activated by ribosome collisions are complex, exhibiting redundancy and competition. We show here that frameshifting is inhibited not only by Mbf1, but also by the ISR components Gcn1 and Gcn20, as well as the NGD regulator Hel2, providing support for a model (Fig. 7A) in which eEF3, Mbf1, Gcn1/ Gcn20 and Hel2 compete to act on the collided ribosome. Competition between the ISR and NGD was previously established based on enhanced induction of the ISR pathway in mutants lacking *HEL2* (Meydan and Guydosh 2020; Yan and Zaher 2021). Here, we find that the effects of eEF3 are held in check by the combined actions of Hel2, Mbf1 and Gcn1/Gcn20, based on the observation that mutation of all three regulators results in a large increase in frameshifting in the *yef3-fs1009* mutant. The argument that frameshifting is prevented by redundancy between Hel2 and Gcn1/Gcn20 functions is supported by the finding that ribosomes also frameshift efficiently in *hel2Δ gcn1Δ* double mutants in which Mbf1 is present. Thus, competition for collided ribosomes as substrates for eEF3, Mbf1, Gcn1/Gcn20 and Hel2 drives the balance of events. These results are consistent with a system with opposing forces in which eEF3 acts to promote frameshifting on collided ribosomes lacking Mbf1, while Mbf1 holds the mRNA and 40S head with Gcn1 and Gcn20 (Sinha et al. 2020; Pochopien et al. 2021) and Hel2 works to remove the stalled ribosome. The redundancy in the three sets of regulators (Gcn1, Gcn20; Mbf1 and Hel2) that all work to restrain frameshifting at the collided ribosome demonstrates the extensive coordination between the translational quality control systems, which allows a plasticity of the response dependent upon the particular problem.

**Figure 7.**
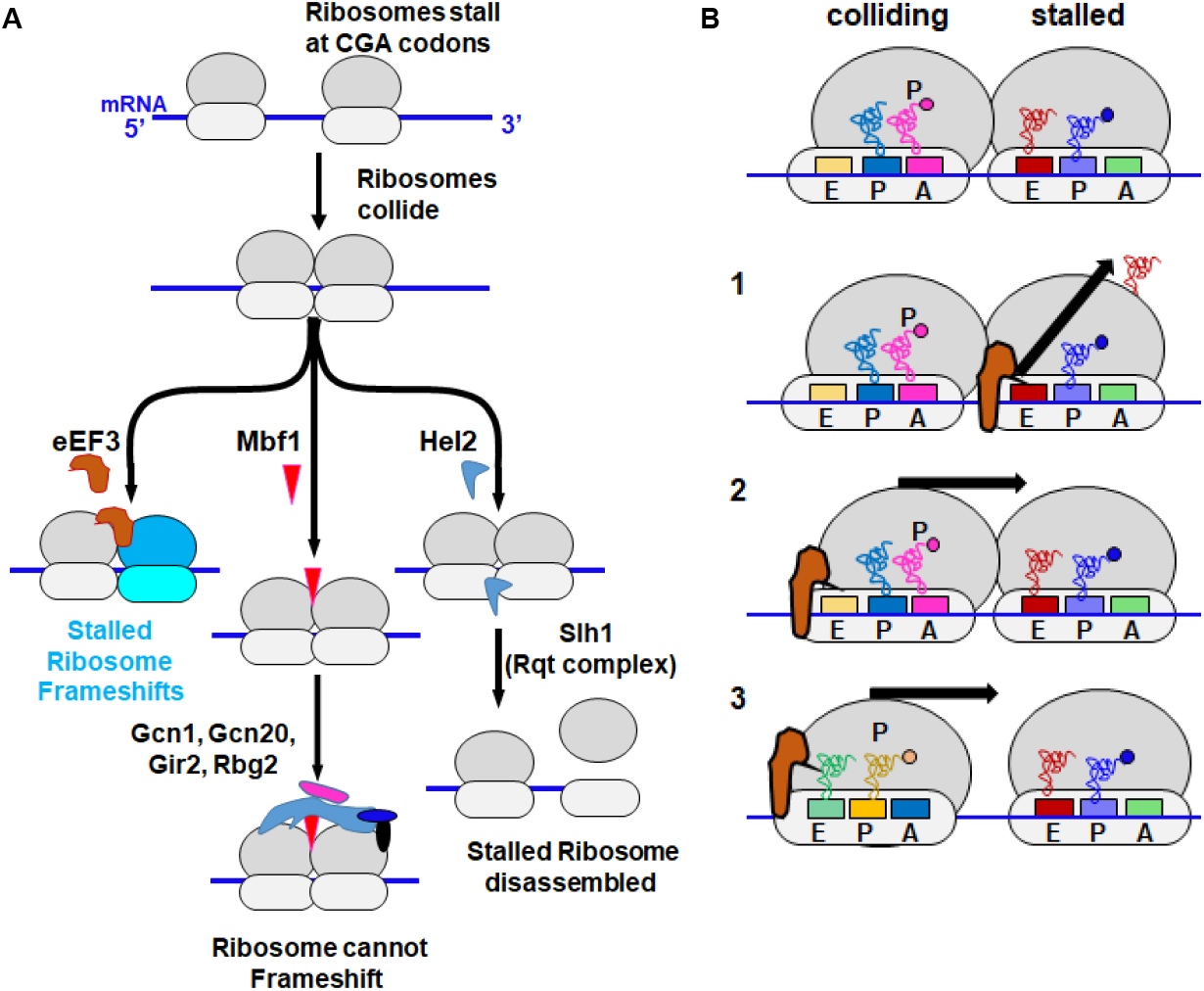
Models of eEF3 functions at collided ribosomes. (A) The competition between eEF3 and other quality control regulators for collided ribosomes determines the outcomes of ribosome collisions. We propose that eEF3 can act on collided ribosomes prior to Mbf1 binding, but that Mbf1 interaction prevents the action of eEF3. Gcn1/Gcn20/Gir2/Rbg2 binding to the Mbf1-bound collided ribosome (Pochopien et al. 2021) likely assists Mbf1 or blocks the interaction of eEF3 due to homologous ribosome binding sites (Marton et al. 1993; Visweswaraiah et al. 2012). Hel2-mediated ubiquitination results in disassembly of the stalled ribosome by Slh1 resulting in depletion of the pool of collided ribosomes. B. Three models for the mechanisms by which eEF3 may induce frameshifting on collided ribosome. Model 1:eEF3 could effect dissociation of the E site tRNA from the stalled ribosome. Model 2: eEF3 could bind the hybrid collided ribosome to finish the translocation reaction into a POST state, which would increase the force on the mRNA. Model 3. eEF3 could be responsible for driving the collided ribosomes into close contact, closing the gap that traps the colliding ribosome in the hybrid state.

The molecular role of eEF3 in translation informs speculation about the likely defect in eEF3 function caused by the *yef3-fs1009* mutation. eEF3 participates in the translocation reaction and in removal of the E site tRNA, based on biochemical, structural and ribosome profiling analysis (Triana-Alonso et al. 1995; Ranjan et al. 2021), and has also been implicated in coupling between exit of the tRNA from the E site and the delivery of tRNA to the A site of the ribosome (Uritani and Miyazaki 1988; Kamath and Chakraburtty 1989; Triana-Alonso et al. 1995; Anand et al. 2003). If the *yef3-fs1009* mutation primarily results in a reduction in the effective amount of functional eEF3 protein, we might expect the mutant to display defects in these activities. The proposal that the mutation affects the amount of functional protein is consistent with observations that the mutation is recessive, confers sensitivity to translational inhibitors, and is at least partially rescued by increased expression of the mutant form of the protein. However, one alternative is that the effects of the *yef3-fs1009* mutation are due to the loss of the conserved C-terminal domain reported to have ribosome binding activity (Kambampati and Chakraburtty 1997); if so, the mutant might primarily affect a specific interaction between eEF3 and collided ribosomes.

We think there are three reasonable models for the proposed role of eEF3 in promoting frameshifting (Fig. 7B). First, eEF3 might work to cause ejection of the E site tRNA from the stalled ribosome, resulting in a ribosome with a single P site tRNA which itself has weak base pairing interactions. In the yeast collided ribosome structure on the CGA-CCG stalling reporter, the stalled ribosome is found in the POST state with tRNAs in the P and E site (Ikeuchi et al. 2019). eEF3 binds tightly to ribosomes in the POST state and promotes the ejection of the E site tRNA (Ranjan et al. 2021). Thus, this model is based on the known activities of eEF3, and it is easy to envision that the extent of frameshifting would depend upon the fraction of stalled ribosomes in which the E site tRNA has been ejected. Second, eEF3 could act on the colliding ribosome to drive it into the leading stalled ribosome and create additional strain on the mRNA. The colliding ribosome has been found in the hybrid state with A/P and P/E tRNAs in an incomplete translocation step and the idea is that the stalled ribosome prevents it from completing translocation (Ikeuchi et al. 2019), but this ribosome, which lacks eEF2, may be an excellent substrate for eEF3 (Ranjan et al. 2021). Third, eEF3 could act on the colliding ribosome prior to or at the point of the actual collision to drive it into closer approximation to the stalled ribosome. We note that Meydan and Guydosh (Meydan and Guydosh 2020) found both 58 and 61 nucleotide disome footprints that differ at their 5’ ends, suggesting different spacing between collided ribosomes. One possibility is that Hel2 and Gcn1 can bind the more widely spaced disomes, but that these disomes do not drive efficient frameshifting because there is less tension on the mRNA in the wider configuration. eEF3, as part of its normal function in elongation, could drive the collided and stalled ribosomes into close approximation (the 58 nucleotides) and this close configuration might drive frameshifting.

We also found evidence that key regulators of the ISR pathway Gcn1 and Gcn20, which bind collided ribosomes with Mbf1 (Pochopien et al. 2021), play a role in reading frame maintenance. To our knowledge, the reading frame maintenance role of Gcn1 and Gcn20 is the only known case in which their function is not tied to ISR induction. The effects of deleting *GCN1* depend on the status of Mbf1 (or uS3): deletion of *GCN1* has relatively small effects on frameshifting when Mbf1 and uS3 are functional, has much greater effects on frameshifting when Mbf1 or uS3 are compromised, but has no additional effect on frameshifting when Mbf1 is removed. Moreover, Gcn1 has no direct contacts with Mbf1, although the architecture of the collided ribosomes containing Gcn1 (and its binding partners) is more compact than that of the collided ribosomes stalled on CGA-CCG (Pochopien et al. 2021). A parsimonious explanation for all of these data is that while Mbf1 is essential to prevent frameshifting, the effects of Gcn1 (and its binding partners) on the overall architecture of the collided ribosome either stabilize Mbf1 or facilitate its function on the collided ribosome. Our initial expectation was that the role of Gcn1/Gcn20 might be to compete directly with eEF3 since there is evidence of such a competition and eEF3 shares extensive homology with ribosome binding domains in Gcn1 and Gcn20 (Marton et al. 1993; Visweswaraiah et al. 2012). However, we did not find the expected increase in frameshifting in the *yef3-fs1009* mutant when *GCN1* or *GCN20* were deleted.

The biological significance of putting Mbf1 into a complex with Gcn1, Gcn20, Rbg2, and Gir2 on the collided ribosome is not immediately obvious. We think it is not likely to be a coincidental pairing, since Mbf1 affects induction of the ISR (Takemaru et al. 1998), and Gcn1 (and Gcn20) affects frameshifting. One idea is that the relationship is used to measure the prevalence of collisions. In that light, we note that Mbf1 is far more abundant in cells (85,474 molecules per cell) than either Gcn1 or Gcn20 (9,432 and 13,281 molecules per cell) (Kulak et al. 2014), an imbalance that could result in reduced ability to prevent frameshifting in cells in which the number of collisions exceeds the capacity of the ISR regulators.

Many aspects of the relationships between the translation machinery and the quality control systems remains to be investigated, including the extent to which different pathways are activated by distinct signals. For example, Yan and Zaher (Yan and Zaher 2021) demonstrated that while ribosome collisions activate both the NGD and ISR pathways, induction of the ISR, but not NGD, is much more efficient with treatments that leave an empty A site in the stalled ribosome. Thus, the NGD and ISR pathways are activated in slightly different ways. One attractive possibility is that frameshifting is driven by a distinct subset of collided ribosomes, perhaps those in which the E site tRNA from the stalled ribosome has been ejected or those in which the collided ribosomes are in close apposition.

## Materials and Methods

### Strains, plasmids, and oligonucleotides

Strains, plasmids, and oligonucleotides used in this study are listed in Supplemental Tables S2-S4. Parents for yeast strains used in this study were BY4741 (*MATa his3Δ leu2Δ0 met15Δ0 ura3Δ0*) or BY4742 (*MATα his3Δ leu2Δ0 lys2Δ0 ura3Δ0)* (Open Biosystems). RNA-ID reporters were constructed as previously described and integrated at the *ADE2* locus, using a *MET15* marker in *MATa* strains and an *S.pombe HIS5* marker in *MATα* strains (Dean & Grayhack, 2012; Gamble et al., 2016; Wolf & Grayhack, 2015; Wang et al., 2018). The *mbf1-R89K* suppressor P15-30 was obtained from YJYW290 (*MATa mbf1-R89K GLN4_(1-99)_-(CGA)_6_+1-URA3*; RFP-*GAL1,10* promoter-*GLN4_(1-99)_-(CGA)_4_+1-*GFP*-MET15 [LEU2 ASC1]*) (Wang et al. 2018).

To obtain Leu^-^ derivatives of P15 and P15-30, cell cultures were grown overnight in YPD, diluted to achieve ∼400 cells/0.1 ml, plated onto YPD, incubated for two days at 30°C and replica plated to SD-Leu and YPD plates. Six Leu^-^ colonies were isolated from each strain, streaked for single colonies on YPD and tested for growth on YPD, YPG, SD-Leu, SD-Ura and SD complete plates at 30°C.

To construct yeast strains in which the 3’ end of the *YEF3* coding sequence was replaced, we assembled integrating plasmids in which base pairs 1640-3135 of the *YEF3* gene (wt or *yef3 G1007V K1009 fs*) extending to +305 in the 3’UTR were fused to a selectable marker (kanR or *K. lactis URA5*), and then followed by 207 base pairs of the *YEF3* 3’ UTR. Plasmids bearing either wild type *YEF3* (pELB1306) or *yef3-fs1009* (pELB1310) coding sequences (nt 1640 through nt 305 of *YEF3* 3’UTR) were fused to a kan^R^ marker in pLB1264, which was derived from pEJYW279, a modified Bluescript vector with a kan^R^ marker (Wang et al. 2018) by cloning 207 bp of *YEF3* 3’UTR (OLB239) into the NheI and NotI sites. The chromosomal *YEF3* gene (*YEF3* wild type or *yef3 G1007V K1009fs*) was PCR amplified (oligos OLB236 and OLB247) and cloned into pELB1264 between XmaI and AatII to create pELB1306 (wt) and pELB1310 (*yef3 G1007V K1009fs*). These plasmids were digested with XmaI and NotI, followed by linear transformation into P15, P15-30, YEVN922 and YLB5853. The *YEF3* gene in the resulting yeast strains was sequence verified.

Similarly, plasmids bearing either wild type *YEF3* (pELB1274) or *yef3-fs1009* (pELB1278) were assembled by fusion to *K.lactis URA5* in pELB1258, which was derived from pECB1330 (a modified Bluescript vector with a *K.lactis URA5* marker) by insertion of the first 207 base pairs of *YEF3* 3’UTR (OLB239) between NotI and NheI. The chromosomal *YEF3* gene (*YEF3* wild type and *yef3 G1007V K1009fs*) was PCR amplified from 1640 base pairs in the *YEF3* coding sequence to 305 bases of 3’ UTR (oligos OLB235 and OLB237) and cloned into ELB1258 between MluI and SacI sites. Following MluI and Not1 digestion, *YEF3* and *yef3-fs1009* were integrated into BY4741 by linear transformation. Both the integrating plasmids and *YEF3* alleles in the resulting Ura+ strains were sequence verified. FOA resistant isolates were selected to obtain strains in which the *K. lactis URA5* marker was removed.

*MBF1 alleles were* introduced into Ura^-^ derivatives of the *YEF3* strains (YLB5691 *YEF3* wt and YLB5715 *yef3 G1007V K1009fs*) by linear transformation of XmaI and NheI digested pEJYW279 (*MBF1-HA*), pEJYW344 (*MBF1-StrepII*) (Wang et al. 2018), and pELB1418 (*mbf1-R89K-HA*). To construct the *mbf1-R89K-HA* plasmid (pELB1418), base pairs 230-411 of the coding sequence from *mbf1-R89K* (OLB256) were cloned into the BamHI and AatII sites of pEJYW279. *YEF3* strains with *MBF1* deletions were constructed by PCR amplification of *mbf1Δ:*kan^R^ with OEVN015 and OJYW125 followed by linear transformation into *YLB5691* and *YLB5715*. The *MBF1* alleles from these strains were verified by sequencing.

The plasmids *YEF3 CEN LEU2* (EEVN250) and *yef3-fs1009 CEN LEU2 (*EEVN246) were constructed in two steps to insert the entire coding sequence of *YEF3* (wt or *yef3 G1007V K1009fs*) with flanking sequences from -714 to +305. In the first step, a gene block (gbEP03) bearing sequences -714 to -652, restriction sites Mlu1 and Xba1, and sequences +245-+305, followed by restriction site AatII was cloned into PstI and EcoRI sites of AVA581 (Alexandrov et al. 2006) to produce EEVN237. The *YEF3* containing plasmids EEVN250 (wt) and EEVN246 (*yef3 G1007V K1009fs*) were constructed using Gibson assembly (Gibson et al. 2009) of the MluI-XbaI digested EEVN237, a PCR product from -714 to base pair 226 in the *YEF3* ORF (using oligonucleotides OEP152 and OEP153 to amplify BY4741 DNA) and Msc1-AatII digested ELB1314 (wt YEF3) or ELB1319 (*yef3 G1007V K1009fs)* to supply the 3286 base pair *YEF3* sequences from 166 in the *YEF3* ORF through +305. Each clone (EEVN250 and EEVN246) was sequence verified.

The *GCN1 CEN LEU2* plasmid EEVN129 was constructed in two steps to insert *GCN1* with flanking sequences (−804 to +341) into the vector AVA581 (Alexandrov et al. 2006). In the first step, a gene block bearing sequences -804 to -342, restriction sites NruI and XmaI, and sequences 7980 in the *GCN1* ORF to +341 in 3’ UTR was cloned into PstI and EcoRI sites of AVA581 (EEVN109). The *GCN1* containing plasmid EEVN129 was obtained by gap repair in yeast, following transformation of BY4741 with NruI and XmaI digested EEVN109, plasmids were isolated using Zymoprep Yeast Plasmid Miniprep II kit, transformed into *E. coli*, isolated by Qiagen minipreps. After verification of the presence of the *GCN1* coding sequence by PCR (OEP063 and OEP064) and restriction digestion, plasmids were tested for functional complementation by transformation into YEVN1004 bearing *gcn1Δ HIS3* and selection on 3-aminotriazole (Hilton et al. 1965; Klopotowski and Wiater 1965). The complementing clone EEVN129 was sequence verified.

Deletions of *MBF1, HEL2, GCN2*, *GCN20, GCN4, YIH1, RBBG2* and *GIR2* were constructed by standard methods using the genomic yeast deletion collection (Giaever et al. 2002) or plasmid cassettes bearing resistance markers (Wach et al. 1994; Goldstein and McCusker 1999; Gueldener et al. 2002).

### Selection for mutants which suppress frameshifting when *MBF1* is defective and identification of mutations

FOA resistant (FOA^R^) mutants were selected from two independent cultures from the P15 strain with the *mbf1-R89K* mutation in YJYW289 (Wang et al. 2018). Strains were grown overnight in three mL YPD at 30°C, harvested, washed twice with sterile water, and re-suspended in 1mL to OD_600_ 0.7. Approximately two million cells were plated on SD-Ura plates containing 50 μg/mL uracil and 500 μg/mL of 5-fluoroorotic Acid (FOA) (Boeke et al. 1987). The selection plates were grown at 30°C for nine days. Single colonies of FOA resistant mutants were streaked onto SD-Ura plates containing 50 μg/mL uracil and 500 μg/mL of FOA. Single colonies from each streak were grown in YP Raf/Gal, spotted onto SD-Ura plates to determine if the mutants displayed an Ura-phenotype and analyzed by flow cytometry to measure GFP and RFP expression. Ura-mutants that had GFP/RFP values less than 60% of the parent, GFP values less than 65% of the parent and RFP values less than 125% of the parent were considered Ura^-^ GFP^-^ mutants, indicative of poor frameshifting efficiency, and were selected for further study.

To identify relevant mutations in *mbf1-R89K* (YJYW290-15) suppressor P15-30, whole genome sequencing was performed on DNA isolated from approximately 30 OD_600_ yeast cells using Lucigen MasterPure Yeast DNA Purification kit (Lucigen catalog: MMPY80200) according to the manufacturer’s directions. Purified DNA (200 μl TE, pH8) was treated with 2 μl RNAse A (10mg/ml stock) at room temperature for one hour, followed by treatment with PCA (Invitrogen 15593-031), then precipitation and washing with ethanol. The pellet was dried and resuspended in 30 μl sterile dH_2_O. Whole genome sequencing was performed by the UR Genomics Research Center.

### Analysis of yeast growth

Appropriate control strains and 2-4 independent isolates of each strain being tested were grown overnight at 30°C in rich media (YPAD or YP Raf/Gal). The strains were diluted in sterile water to obtain 0.5 OD_600_ (for four spot tests) or 0.05 OD_600_ (for three spot tests) followed by 10-fold serial dilutions in sterile water. Two μl of diluted cells were spotted onto the indicated plates and grown at various temperatures for a minimum of two days.

### Western blotting

Cells from 100 mL YP Raf/Gal culture were grown to an OD_600_ of 0.8-1.2, harvested by centrifugation and resuspended in 120 μL – 160 μL extraction buffer (20mM Hepes pH 7.5, 1M NaCl, 5% Glycerol, 2mM 2-mercaptoethanol (BME) 1mM pefabloc, 2.5 μg/mL leupeptin and 2.5 μg/ml pepstatin) (Alexandrov et al. 2004) and 0.5 mm Zirconia /Silca beads (BioSpec #11079105z) and lysed with vortex (5 repeats of 1 minute vortex followed by 1 minute in ethanol-ice) essentially as described previously (Gelperin et al. 2005). The cell lysates were collected by centrifugation at 4°C for 10 minutes at 13,000 RPM. The crude extracts were separate by SDS PAGE on 4-20% Criterion TGX precast midi protein gels (BioRad #5671094) and transferred to a 0.2 μm nitrocellulose membrane (BioRad #1620112) and blotted as described previously (Gelperin et al. 2005). eEF3 protein was detected with anti-eEF3 antibody (Kerafast ED7003) and Glucose-6-phosphate dehydrogenase (G-6-PDH) with anti-G-6-PDH antibody (Sigma A9521). Blots were probed with IgG Goat anti-Rabbit (BioRad 170-6515) for anti-eEF3 and anti-G-6-PDH secondary antibodies and developed with Pierce ECL plus western blotting substrate kit (Thermo Scientific 32132).

### Coomassie stained gel

Crude extracts of the given strains were separated on 4-20% Criterion TGX precast midi protein gels (BioRad #5671094). The gel was washed in fixing solution (40% Ethanol 10% Acetic Acid) for 15 minutes and rinsed in deionized water three times for 5 minutes each. The gel was stained in QC colloidal Coomassie stain (BioRad #1610803) for 17-20 hours, followed by de-staining in deionized water for three hours, changing the water every hour.

### Flow Cytometry

Yeast strains containing modified RNA-ID reporters were grown at least 24 hours prior to analysis at 30°C in YP media (for strains without a plasmid) or appropriate synthetic drop-out media (for strains with a plasmid) containing 2% raffinose + 2% galactose + 80 mg/L Ade. The cell culture was diluted six hours before analysis such that the culture had a final OD_600_ between 0.8-1.1. Analytical flow cytometry and downstream analysis were performed for four to six independent isolates of each strain (Outliers were rejected using a Q test with >90% confidence level) as previously described (Dean and Grayhack 2012). P values were calculated using a one-tailed or two-tailed homoscedastic or heteroscedastic t test in Excel, as indicated in the source data for relevant figures.

### RT-qPCR

mRNA measurements with reverse transcription (RT) and quantitative PCR were performed as described previously (Gamble et al. 2016)

### Plasmid transformation

Yeast strains bearing plasmids were transformed as previously described (Schiestl and Gietz 1989).

### Linear Transformation

Yeast strains bearing RNA-ID reporters and chromosomal deletions were obtained by linear transformation as previously described (Gietz and Woods 2002).

### Alignment

Amino acid sequence alignments were obtained using - multAlin (Corpet 1988) http://multalin.toulouse.inra.fr/multalin/

## SUPPLEMENTAL MATERIAL

Supplemental material is available for this article.

## ACKNOWLEDGEMENTS

We thank Eric Phizicky, Christina Brule, Thareendra De Zoysa, Dmitri Ermolenko, Alayna Hauke, Erin Marcus, Elaine Sia, Monika Tasak, and Yi-Tao Yu for discussions of the science, and Eric Phizicky for comments on the manuscript. We thank the University of Rochester Genomics Research Center for high-throughput sequencing, including library construction, sequencing, and primary data analysis for this study, and the URMC Flow Cytometry Resource for technical support. This work was supported by NIH grant R01 GM118386 to E.J.G. L.B.H. was also supported by a NIH T32 Training Grant in Cellular, Biochemical and Molecular Sciences (GM68411).

---

**Supplemental Table S1.**
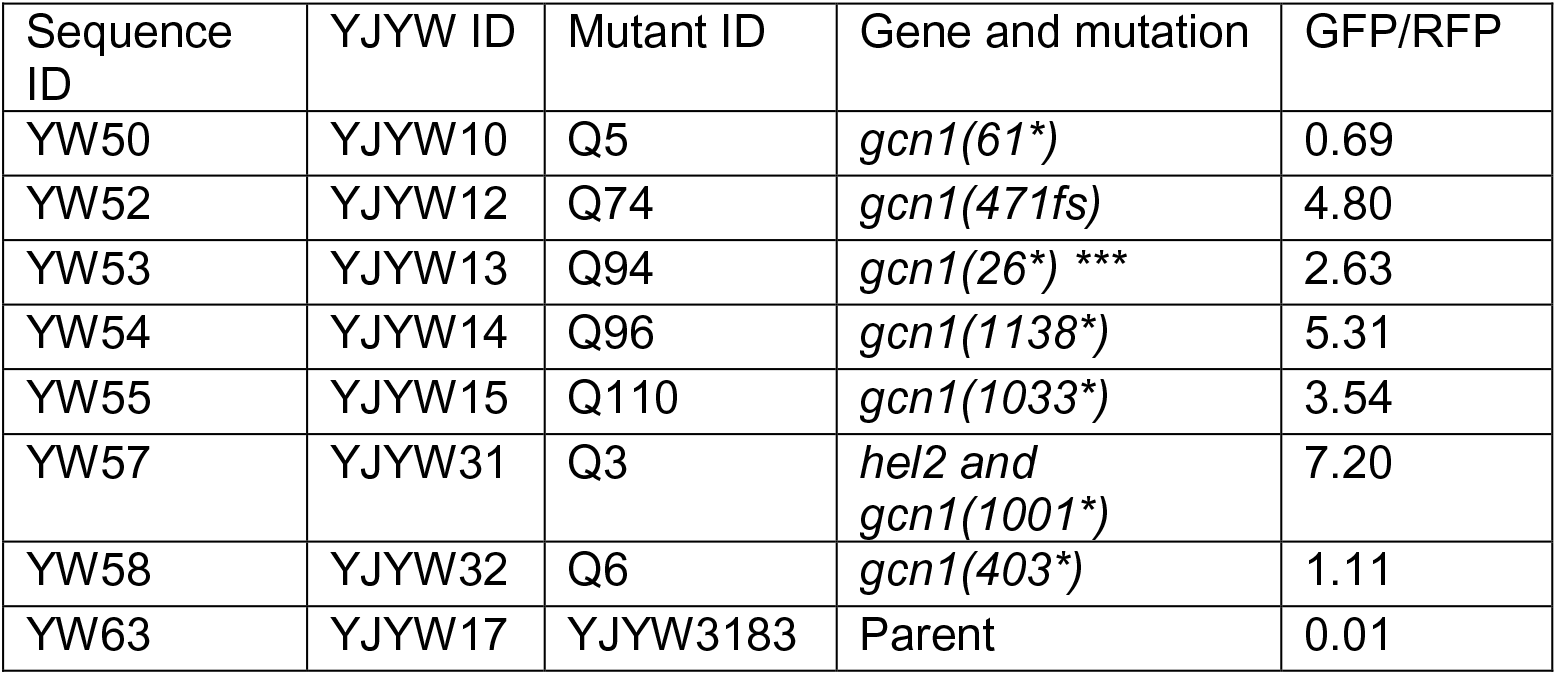

| Sequence ID | YJYW ID | Mutant ID | Gene and mutation | GFP/RFP |
| --- | --- | --- | --- | --- |
| YW50 | YJYW10 | Q5 | *gcn1*(61*) | 0.69 |
| YW52 | YJYW12 | Q74 | *gcn1*(471fs) | 4.80 |
| YW53 | YJYW13 | Q94 | *gcn1*(26*) *** | 2.63 |
| YW54 | YJYW14 | Q96 | *gcn1*(1138*) | 5.31 |
| YW55 | YJYW15 | Q110 | *gcn1*(1033*) | 3.54 |
| YW57 | YJYW31 | Q3 | *hel2* and<br>*gcn1*(1001*) | 7.20 |
| YW58 | YJYW32 | Q6 | *gcn1*(403*) | 1.11 |
| YW63 | YJYW17 | YJYW3183 | Parent | 0.01 |

**Supplemental Figure S1. Suppressor P15-30 is recessive.** Expression of frameshifted GFP/RFP is significantly lower in the haploid *MAT**a*** P15-30 compared to that in *MAT**a*** P15, but expression of the frameshifted GFP/RFP is nearly identical for diploids of both *MAT**a*** P15 and *MAT**a*** P15-30 mated with *MATα mbf1Δ* (*MAT**a***/*MATα mbf1Δ*/*mbf1-R89K YEF3/YEF3 and MAT**a***/*MATα mbf1Δ*/*mbf1 R89K YEF3/yef3 G1007V K1009fs*).

**Supplemental Figure S2. Amino acid sequence alignment of eEF3 from six *Ascomycete* fungi across several different clades and the *yef3-fs1009* mutant using MultAlin** (http://multalin.toulouse.inra.fr/multalin/) (Corpet 1988). The color text represents the level of consensus for each residue (Blue: 50-90%, Red: >90%).

**Supplemental Figure S3. The *yef3-fs1009* mutation alters eEF3 amounts, which can be restored by additional copies of the mutant form, and does not affect TY1 frameshifting.** (A) Strains with the *yef3-fs1009* mutation have reduced levels and altered migration of eEF3 protein compared to otherwise isogenic strains with wild type *YEF3*. Crude extracts from the indicated strains were subjected to SDS-PAGE and Coomassie-staining. (B) eEF3 levels in the *yef3-fs1009* mutant expressing *yef3-fs1009* on a *CEN* plasmid exceed the levels of eEF3 in a *YEF3* wild type strain with an empty vector. Crude lysates separated by SDS-PAGE were subjected to Western analysis with anti-eEF3 and anti-glucose-6-phosphate-dehydrogenase (G6PD) antibodies. (C) The *yef3-fs1009* mutant does not affect *TY1* programmed frameshifting. GFP/RFP levels from GFP constructs bearing the TY1 frameshifting sequence in the indicated reading frames were examined in strains with *YEF3* wild type and *yef3-fs1009*.

**Supplemental Figure S4. CGA codon pairs are inhibitory in the *yef3-fs1009* mutant with either wild type *MBF1* (A) or completely lacking *MBF1 (mbf1Δ*) (B).** GFP/RFP protein (fluorescence), mRNA and protein/mRNA were examined from the inhibitory (CGA-CGA)_3_ and optimal (AGA-AGA)_3_ in-frame reporters as well as the frameshifted (CGA-CGA)_3_+1 reporter. As expected, the GFP/RFP from the frameshifting reporter (CGA-CGA)_3_+1 was near background levels with wild type MBF1(A), but was much greater in *mbf1Δ* strains (B). Frameshifted GFP/RFP was significantly lower in the *yef3-fs1009 mbf1Δ* mutant compared to the *YEF3 mbf1Δ* mutant.

**Supplemental Figure S5. Effects of *GCN1*, *RPS3* and *GCN20* on frameshifting depend upon the mutations in these genes, are not mediated through effects on in-frame expression and do not depend upon a functional copy of *GCN2*.** (A) Schematic of the parental strain YYJYW290 used for selection of mutants that promote frameshifting (Wang et al. 2018). (B) Exogenous expression of wild type *RPS3* or frameshifting mutant *RPS3-K108E* results in expected effects on frameshifted GFP/RFP. Wild type *RPS3* suppresses the frameshifting in the *RPS3-S104Y* single mutant and in the *gcn1Δ RPS3-S104Y* double mutant but has no effect on frameshifting in the *gcn1Δ* single mutant. *RPS3-K108N* results in increased frameshifting in the wild type, *gcn1Δ*, and *RPS3-S104Y* mutants, but has no detectable effect in the *gcn1Δ RPS3-S104Y* double mutant. (C) Expression of the in-frame reporters is not affected by the *gcn1Δ* mutation*, RPS3S104Y* mutation, or the *gcn1Δ RPS3 S104Y* double mutations. (D) Expression of *GCN20* results in reduced frameshifting in the *gcn20Δ RPS3 S104Y* double mutant but has no effect on frameshifting in the *gcn1Δ RPS3 S104Y* double mutant. (E) The effects of *gcn1Δ* on frameshifting in the *RPS3 S104Y* mutant do not require Gcn2 function. Expression of the frameshifted reporter in *gcn1Δ RPS3 S104Y* mutants is not affected by deletion of *GCN2*.

**Supplemental Figure S6. Exogenous expression of *HEL2* and *MBF1* modulate frameshifting.** (A) Expression of *HEL2* complements *hel2Δ* and *hel2Δ gcn1Δ* in both *YEF3* and *yef3-fs1009* strains, resulting in a reduction in frameshifted GFP/RFP to levels at or below those in the corresponding *HEL2* strain. Moreover, expression of *HEL2* also results in a reduction in frameshifted GFP/RFP in *gcn1Δ* mutants. (B) Expression of *MBF1* from a multicopy plasmid resulted in a reduction in frameshifted GFP/RFP in all strains, suppressing frameshifting to near background levels in the parent and single mutant strains. However, MBF1 expression only partially suppressed frameshifting in the *hel2Δ gcn1Δ* mutants.

